# Influenza A virus-induced PI4P production at the endoplasmic reticulum involves ATG16L1 and promotes the egress of viral ribonucleoproteins

**DOI:** 10.1101/2024.11.29.625996

**Authors:** Carla Alemany, Juliane Da Graça, Quentin Giai-Gianetto, Sylvain Paisant, Maud Dupont, Thibaut Douché, Catherine Isel, Cédric Delevoye, Lydia Danglot, Mariette Matondo, Etienne Morel, Jean-Baptiste Brault, Nadia Naffakh

## Abstract

The genomic RNAs of influenza A viruses (IAVs) are replicated in the nucleus of infected cells in the form of viral ribonucleoproteins (vRNP) before being exported to the cytoplasm. The small GTPase RAB11A is involved in the transport of vRNPs to the sites of viral assembly at the plasma membrane, but the molecular mechanisms involved remain largely unknown. Here we show that IAV infection remodels the architecture of the endoplasmic reticulum (ER) sheets, where vRNPs tend to accumulate in the absence of RAB11A. To decipher the interplay between RAB11A, vRNPs and the ER, we investigated viral-induced perturbations of RAB11A proximity interactome. To this end, we generated cells stably expressing a TurboID-RAB11A fusion protein and performed biotin-based proximity labeling upon viral infection. We found that cellular regulators of phophatidylinositol-4-phosphate (PI4P) homeostasis, including the autophagic and stress response protein ATG16L1, are significantly enriched at the vicinity of RAB11A in infected cells. Infection induces an increase in cellular PI4P levels in an ATG16L1-dependent manner, while ATG16L1 relocalizes to ER membranes upon infection. Depletion of ATG16L1 decreases the co-distribution of vRNPs with PI4P punctae on ER membranes, and reduces the accumulation of vRNPs at the plasma membrane as well as the production of IAV infectious particles. Our data extend to IAVs the notion that viruses can modulate the metabolism and localization of phosphoinositides to control host membrane dynamics and point to the ER as an essential platform for vRNP transport. They provide evidence for a pivotal role of ATG16L1 in regulating the identity of endomembranes and coordinating RAB11A and PI4P-enriched membranes to ensure delivery of vRNPs to the plasma membrane.

## Introduction

Influenza A viruses (IAVs) present continuous animal and public health challenges. Human-adapted IAVs recur every year due to antigenic variation, as seasonal IAVs [1]. Wild aquatic birds and domestic species are hosts to a dynamic pool of IAVs, which are responsible for epizootic, zoonotic and potentially pandemic outbreaks [2]. Sporadically, as a consequence of the segmentation of their genome into a bundle made of 8 distinct viral RNAs (vRNAs), novel and possibly devastating IAVs are generated through co-infection and genetic mixing of vRNAs from animal and human IAVs. The molecular mechanisms of vRNA intracellular transport from the nucleus to the plasma membrane and vRNA assembly into bundles, which are critical for reassortment, are only partially understood [3]. Elucidating this aspect of the viral cycle will help in the broader goal to achieve better prevention and treatment of the disease.

Each influenza vRNA, a single-stranded RNA of negative polarity, is associated with viral proteins into macromolecular complexes called viral ribonucleoproteins (vRNPs) that are 30-120 nm in length and 12-15 nm in diameter. Within each vRNP, the vRNA adopts a closed conformation by base-pairing of the 3’ and 5’ ends. The resulting ∼15 bp-long duplex associates with one copy of the viral polymerase, a PB1-PB2-PA heterotrimer, while the remaining RNA is bound by the nucleoprotein (NP). Upon viral entry, vRNPs are released in the cytoplasm and transported into the nucleus, where they serve as templates for transcription and replication of the viral genome [4]. Neo-synthesized vRNPs may become templates for new rounds of transcription/replication, or exit the nucleus through the CRM1-dependent nuclear export pathway. They are then transported towards the plasma membrane where they get assembled with other viral proteins and glycoproteins, enabling the budding of new virions [3].

There is much evidence that the RAB11A small GTPase is involved in vRNP trafficking and assembly of IAV genome, including the fact that it interacts directly with the PB2 component of vRNPs (reviewed in [5]). At 4-6 hours post-infection, vRNPs start to accumulate in a perinuclear region close to the microtubule organizing center (MTOC), where they colocalize with RAB11A-positive membranes. At later time points in infection, inclusions containing vRNPs and RAB11A can be observed throughout the cytoplasm [6]. RAB11A is a key player of (and used as a marker for) recycling endosomes and its best-documented function is to regulate the recycling of internalized proteins, from endosomes to the plasma membrane [7]. Therefore, it was initially proposed that RAB11A mediates the docking of vRNPs to recycling endosomes in the vicinity of the MTOC, after which recycling endosomes carry vRNPs towards the plasma membrane along microtubules. However, this model is now called into question by the fact infected cells show i) alterations in the efficiency of RAB11A recycling pathway, very likely due to the fact vRNPs hinder RAB11A binding to RAB11-FIP effectors [8,9], and ii) alterations in the intracellular distribution of RAB11A, which is found in close proximity with endoplasmic reticulum (ER) membranes [10,11]. It was proposed that the disruption of RAB11A canonical function leads to the concentration of recycling endosomes coated with vRNPs condensates as a mechanism to facilitate vRNP bundling and assembly of the viral genome [9,10].

Alongside recycling endosomes, the ER compartment is also altered in IAV-infected cells. We previously reported that IAV infection induces a progressive remodeling and tubulation of ER membranes around the MTOC and all throughout the cell [11]. RAB11A and vRNPs were detected close to the ER membranes and at the surface of a new type of RAB11A-positive vesicles, distinct of recycling endosomes, which were named irregularly coated vesicles (ICVs). Some ICVs are observed very close to the ER or the plasma membrane, suggesting that these vesicles could be transporting vRNPs between the two membrane compartments. It was recently showed that the viral hemagglutinin (HA), a glycoprotein which travels through the ER-Golgi secretory pathway, drives the formation of remodeled membrane compartments on which vRNPs tend to accumulate [12]. Interestingly, expression of a dominant-negative mutant of RAB11A decreases vRNP association with HA-remodeled membranes [12] as well as the formation of ICVs [11].

Many questions remain to be answered, notably regarding the site(s) at which RAB11A and vRNPs cooperate, the biogenesis of vRNP-coated vesicles, and their subsequent transport to virion assembly sites at the plasma membrane. Here, we aimed at unraveling the interplay between RAB11A, the ER membranes and vRNPs. Using a combination of confocal and STED microscopy, we extend our previous observations and show that IAV infection specifically alters the distribution of ER sheets. We find that in RAB11A-depleted cells, vRNPs tend to accumulate at the vicinity of remodeled ER membranes. To decipher the interplay between RAB11A, vRNPs and the ER, we combined proximity-labeling and affinity purification-mass spectrometry on cells stably expressing a TurboID-RAB11A fusion protein. We found that several cellular regulators of phosphoinositide (PI) homeostasis, including the autophagy-related protein ATG16L1 [13], are enriched in the proximity-interactome of RAB11A in IAV-infected compared to uninfected cells.

Phosphoinositides (PI) represent a minor fraction of total cellular phospholipids, yet they play key roles in mediating membrane signal transduction, shaping and defining the membrane identity of organelles, and regulating vesicular trafficking (reviewed in [14,15]). PI are produced from a phosphatidylinositol precursor, the inositol ring of which can be phosphorylated at the D3, D4 and D5 positions. As a result, distinct phosphaditylinositol monophosphates (PI3P, PI4P and PI5P), phosphaditylinositol biphosphates (PI(3,4)P_2_, PI(3,4)P_2_, PI(3,4)P_2_) and one phosphaditylinositol triphosphate (PIP_3_) can be produced. Multiple kinases, phosphatases and regulatory proteins are controlling the spatial and temporal turnover of PIs, and the specific composition and dynamics of PI pools at specific endomembrane sites. The cross-talk between PIs and RAB GTPases plays a major role in maintaining this “membrane code” and controlling membrane remodeling and trafficking [16].

Several positive-sense RNA viruses manipulate the PI regulatory machinery and hijack phosphaditylinositol-4 kinases (PI4K) to induce the formation of modified, PI4P-enriched endomembrane compartments referred to as viral replication organelles (reviewed in [17]). However, it remained unknown whether PI metabolism is rewired during IAV infection and whether it plays a role in vRNP trafficking. Here we show that the PI3P/PI4P balance is altered and the pool of PI4P associated with the ER increases upon IAV-infection, through a pathway that involves ATG16L1. Depletion of ATG16L1 affects the co-distribution of PI4P punctae with vRNPs on ER membranes, delays the accumulation of vRNPs at the plasma membrane, and decreases the production of IAV infectious particles. Hence, we propose a working model in which the vRNPs get associated to a remodeled ER, where ATG16L1 mediates, together with RAB11A, a local production of PI4P, and thereby promotes the delivery of vRNP-coated vesicles to the plasma membrane.

## Results

### IAV infection specifically remodels the ER architecture

We sought to extend previous evidence that the ER compartment is altered in IAV-infected cells. To this end we performed confocal microscopy using antibodies specific for the cytoskeleton-linking membrane protein 63 (CLIMP63) and the reticulon 3 protein (RTN3), known to be enriched in ER sheets and ER tubules, respectively [18,19]. A549 cells were infected at a high multiplicity of infection with the A/WSN/33 (WSN) or A/Victoria/3/75 (VIC) virus, fixed at 8 hours post-infection (hpi) and stained for the endogenous RTN3 and CLIMP63 proteins, and the viral NP. The NP staining is both nuclear and cytoplasmic, as expected during the late phase of the viral life cycle. The RTN3 staining shows no significant alteration in IAV-infected compared to mock-infected cells (**Fig 1A**). In contrast the CLIMP63 staining is markedly altered upon IAV infection, exhibiting an irregular and very extended distribution compared to the dense and homogeneous perinuclear signal in mock-infected cells.

**Legend of Fig 1.**
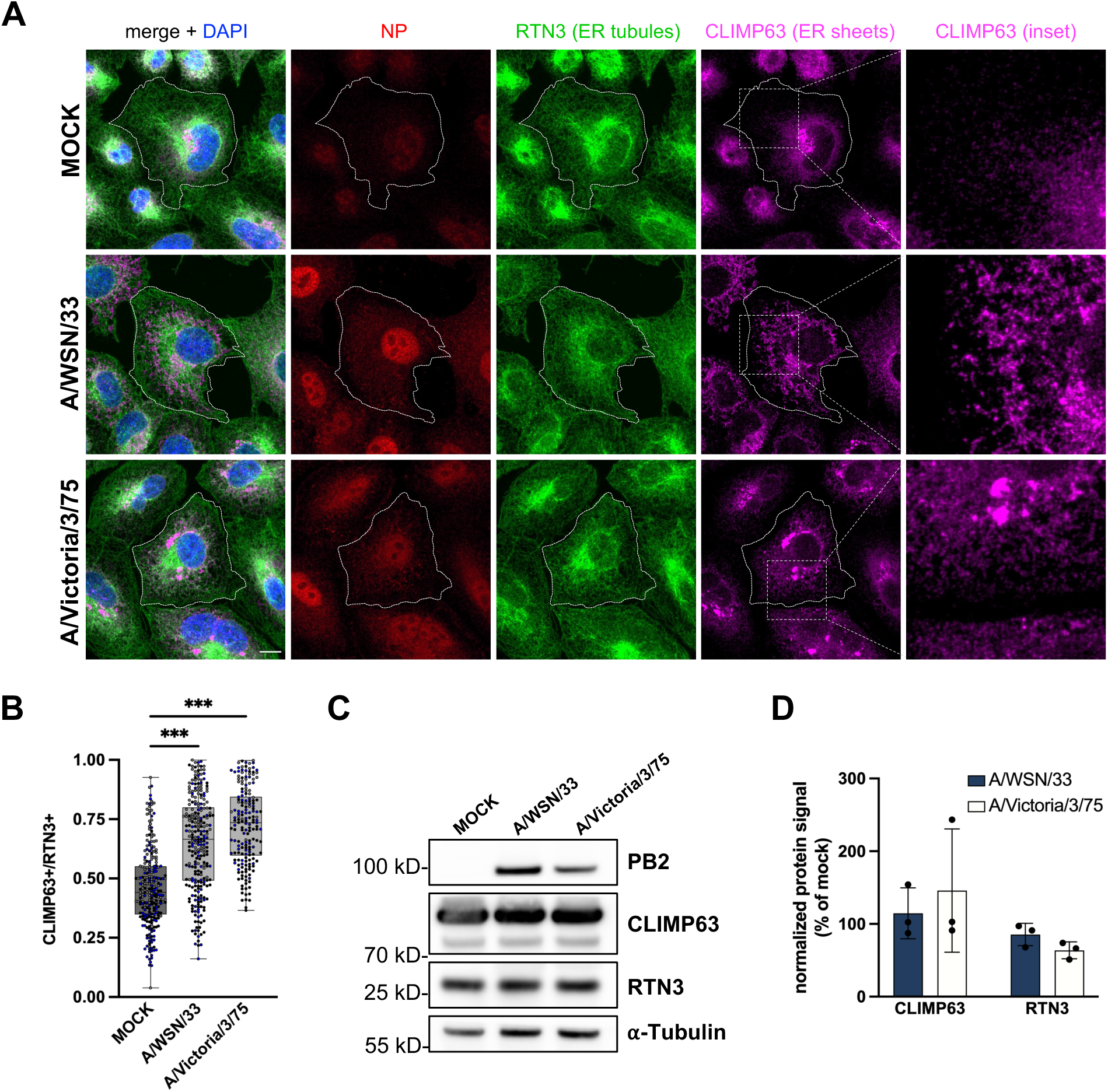
IAV infection specifically remodels the ER sheets. **A.** A549 cells were infected with WSN or VIC at a MOI of 5 PFU/cell for 8 h, or mock-infected. Fixed cells were stained for the viral NP and the cellular markers for ER sheets and ER tubules, CLIMP63 and RTN3, respectively. The RTN3 staining was used to delineate the cell edges. Nuclei were stained with DAPI and cells were imaged with a confocal microscope. Scale bar: 10 µm. **B.** A549 cells treated as in A were analyzed with the Cell Profiler software in order to segment CLIMP63+ and RTN3+ areas based on the Otsu thresholding method. The ratios of the CLIMP63+ area to the RTN3+ area are shown. Each dot represents one cell, and the data from three independent experiments are shown (black, grey and white dots). The median and interquartile values are represented as box-plots (200-350 cells per condition). **\*\*\*** : p-value < 0.001, one-way ANOVA. **C-D.** A549 cells were infected or mock-infected as in A. At 8 hpi total cell lysates were prepared and analysed by western blot, using the indicated antibodies. (C) Cropped blots of one representative experiment out of three are shown. (D) The signals for CLIMP63 and RTN3 are normalised over the tubulin signal and expressed as percentages (100% : mock-infected cells). The data shown are the mean ± SD of three independent experiments. No significant difference is detected between infected and mock-infected cells (two-way ANOVA with Sidak’s multiple comparison test).

We quantified the IAV-induced remodeling of ER sheets using the cell image analysis software Cell Profiler [20,21]. Cells were segmented into an RTN3+ area, used as a proxy for the total cell area (delineated by a dotted line in **Fig 1A**), and another CLIMP63+ area. The ratio of the CLIMP63+ area over the RTN3+ area is significantly higher in WSN-and VIC-infected cells (median values of 0.65 ± 0.20 and 0.72 ± 0.16, respectively) than in mock-infected cells (0.46 ± 0.17, p<0.001) (**Fig 1B**). In the same conditions of infection, the steady-state levels of CLIMP63, as assessed by western blot on total cell lysates, remain unchanged (**Fig 1C-D**). Therefore, the increase in the CLIMP63+ to RTN3+ area ratio is indicative of a redistribution rather than an overproduction or overaccumulation of CLIMP63. The staining for CLIMP63 and the viral HA surface glycoprotein, which travels through the ER-Golgi pathway, largely overlap, indicating that CLIMP63 remains located within the ER (**S1A-B Fig**). In cells infected with Zika virus (ZIKV), a flavivirus that remodels the ER to generate cytosolic viral replication factories, a very distinct pattern is observed with both RTN3 and CLIMP63 being relocalized and concentrated in the same area as the NS3 viral antigen, a marker for viral factories (**S1C Fig**). In summary, our data demonstrate that IAV induces a specific alteration of ER sheet architecture, characterized by an extension towards the cell periphery.

### vRNPs accumulate at the vicinity of ER membranes upon RAB11A depletion

RAB11A depletion leads to an alteration of the late stages of IAV infection, notably the distribution pattern of vRNPs in the cytoplasm, a delayed delivery at the plasma membrane, and a reduction in the production of infectious viral particles [6,22–24]. This points to a role for RAB11A in vRNP transport, but its exact mechanism of action and how it is related to the ER remodeling remain unclear. Here we assessed the impact of RAB11A depletion on the remodeling of CLIMP63+ ER membranes and their proximity with vRNPs. As the NP staining largely coincides with PB2 protein (**S2A-B Fig**), NP signal was used as a proxy to visualise vRNPs in the cytoplasm. Prior to infection, A549 cells were treated with RAB11A-specific siRNAs or Non-Target (NT) siRNAs, and RAB11A knock-down efficiency was confirmed by western blot analysis (**S5A Fig**) and by immunofluorescence (**Fig 2A**). The depletion of RAB11A prevents the formation of large cytoplasmic inclusions of vRNPs upon viral infection (**Fig 2A**), as expected from previous studies [6,22], but does not affect the remodeling of CLIMP63+ membranes (**Fig 2A** and **S2C Fig).** Strikingly, in RAB11A-depleted cells the viral NP signal is found at the vicinity of CLIMP63+ membranes in the perinuclear region (**Fig 2A-B**). While a similar distribution of NP in a reticulated pattern in the perinuclear region of RAB11A-depleted cells has already been reported by Eisfeld et al [6], our observations suggest that the sites where vRNPs accumulate in the absence of RAB11A correspond to ER membranes.

**Legend of Fig 2.**
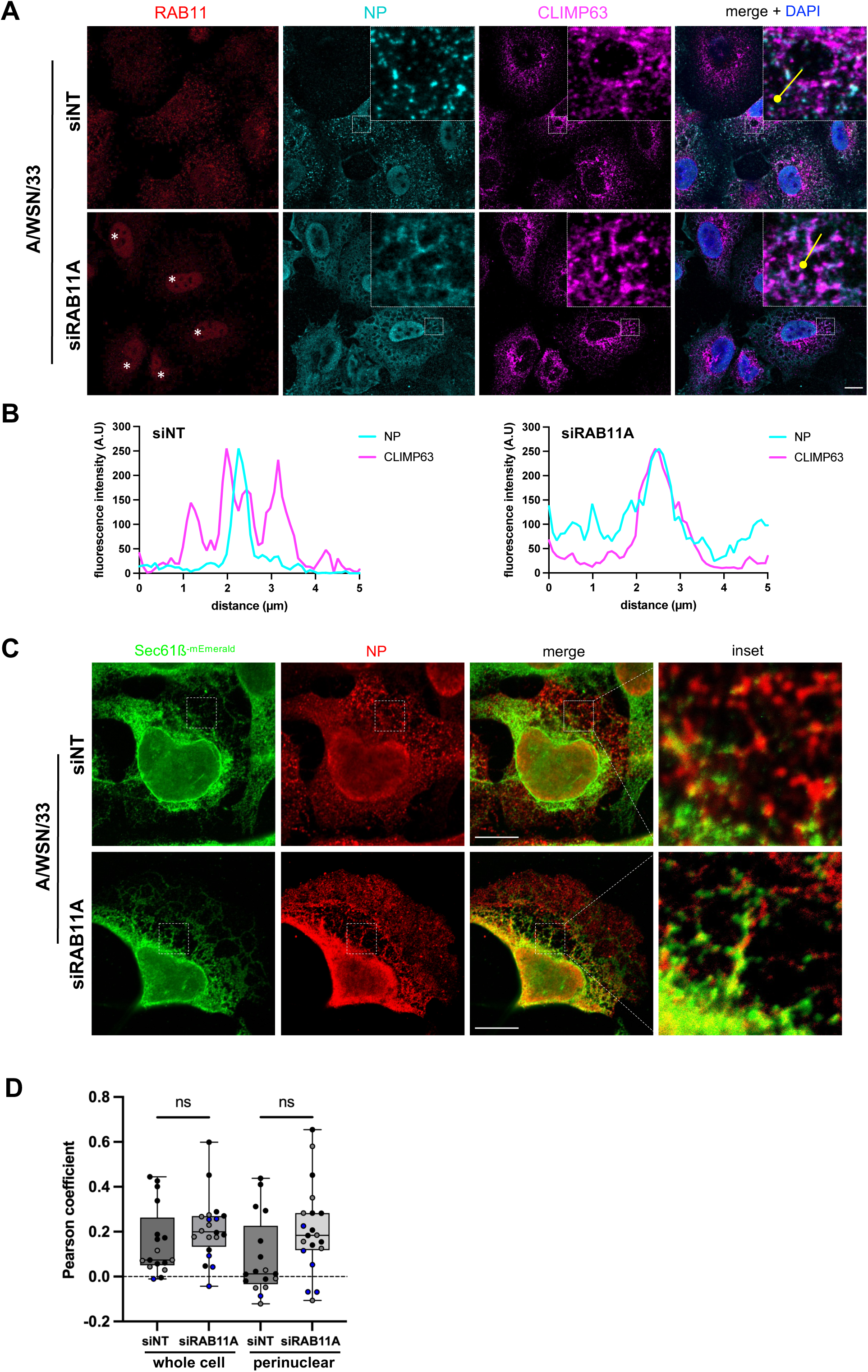
Impact of RAB11A depletion on ER remodeling and vRNP localisation upon infection with the WSN virus. **A.** A549 cells were treated with RAB11A-specific or control Non-Target (NT) siRNAs for 48 h, and subsequently infected with WSN at a MOI of 5 PFU/cell for 4 h. Fixed cells were stained for the viral NP and the cellular RAB11 and CLIMP63 proteins. Nuclei were stained with DAPI and cells were imaged with a confocal microscope. White stars: non-specific nuclear staining enhanced upon siRNA treatment. Scale bar: 10µm. **B.** Fluorescence intensity profiles for NP (red) and CLIMP63 (magenta) along the white lines drawn in panel A (merge insets), starting from the knob. **C.** U2OS cells stably expressing the ER translocon Sec61ß fused to mEmerald (U2OS-Sec61ß-mEmerald) were treated with RAB11A-specific or control Non-Target (NT) siRNAs for 48 h, and subsequently infected with WSN at a MOI of 5 PFU/cell for 8 h. Fixed cells were stained for NP and Sec61ß-mEmerald (anti-GFP antibody) and images were acquired using STED microscopy. Scale bar: 10µm. **D.** U2OS-Sec61ß-mEmerald cells treated as in A were analyzed to assess colocalization of NP and Sec61ß, using a pixel-based method to determine the Pearson coefficient on a region of interest corresponding to the whole cell or to the perinuclear region, as defined in the Methods section. Each dot represents one cell, and the data from three independent experiments are shown (black, grey and white dots. The median and interquartile values are represented as box-plots (18 and 19 cells treated with the NT or RAB11A-specific siRNAs, respectively). ns: non-significant (unpaired t-test).

To further document this point, we used Stimulated Emission Depletion Microscopy (STED), which overcomes the diffraction-limited resolution of confocal microscopes and enables super-resolution imaging [25]. To this end, we used a U20S cell line characterized by a large and flat cytoplasmic area, stably expressing the ER protein Sec61ß fused to mEmerald, and therefore particularly convenient for imaging of the ER [26]. Overexpressed Sec61ß was shown to label both the ER sheets and tubules [18]. This method allowed us to confirm that in RAB11A-depleted cells, vRNPs can be observed along membrane structures in the immediate vicinity of Sec61ß-positive membranes (**Fig 2C**). The Pearson correlation coefficient was used to assess the covariance of Sec61ß and NP signal levels in control and RAB11A-depleted cells. We computed the Pearson coefficient without any thresholding, therefore taking into consideration both the diffuse and aggregated NP signal. The data showed low Pearson coefficient values, which is consistent with the fact that only a fraction of the NP signal is observed at the vicinity of ER membranes, with the NP and Sec61ß signals showing partial overlap. Interestingly however, we observed a non-significant trend to increase in RAB11A-depleted cells (0.15 and 0.21 in siNT- and siRAB11A-treated cells, respectively), this trend being more pronounced in the perinuclear region (0.08 and 0.21 in siNT- and siRAB11A-treated cells, respectively) (**Fig 2D**).

Taken together, our data strongly suggest that that after exiting the nucleus, vRNPs are targeted to ER membranes independently of RAB11A.

### The proximity interactome of RAB11A is enriched in ATG16L1 and phosphoinositide-metabolizing enzymes upon IAV infection

To decipher the interplay between RAB11A, vRNPs and the ER, we determined to what extent its protein interactome is modified upon IAV infection. To this end, we used a TurboID-based proximity labeling approach, most suitable for the detection of weak or transient protein-protein interactions (**Fig 3A**). TurboID is a highly active biotin ligase, it rapidly converts biotin into biotin–AMP, a reactive intermediate which covalently labels neighbouring proteins within a 10-30 nm labeling radius [27]. The RAB11A protein fused to TurboID was stably expressed in A549 cells by lentiviral transduction, and we isolated a clonal population of A549-TurboID-RAB11A cells showing comparable steady-state levels for the recombinant TurboID-RAB11A and the endogenous RAB11A proteins, as assessed by western-blot (**S3A Fig**). Upon immunostaining, the TurboID-RAB11A protein showed the same subcellular distribution as the endogenous RAB11A, *i.e*. regular cytoplasmic punctae concentrated in a perinuclear area in uninfected cells, larger and irregular cytoplasmic punctae strongly colocalizing with the viral NP in infected cells (**S3B Fig**). Addition of biotin to the culture medium during 10 minutes led to a detectable accumulation of biotinylated proteins (**S3C Fig**).

**Legend of Fig 3.**
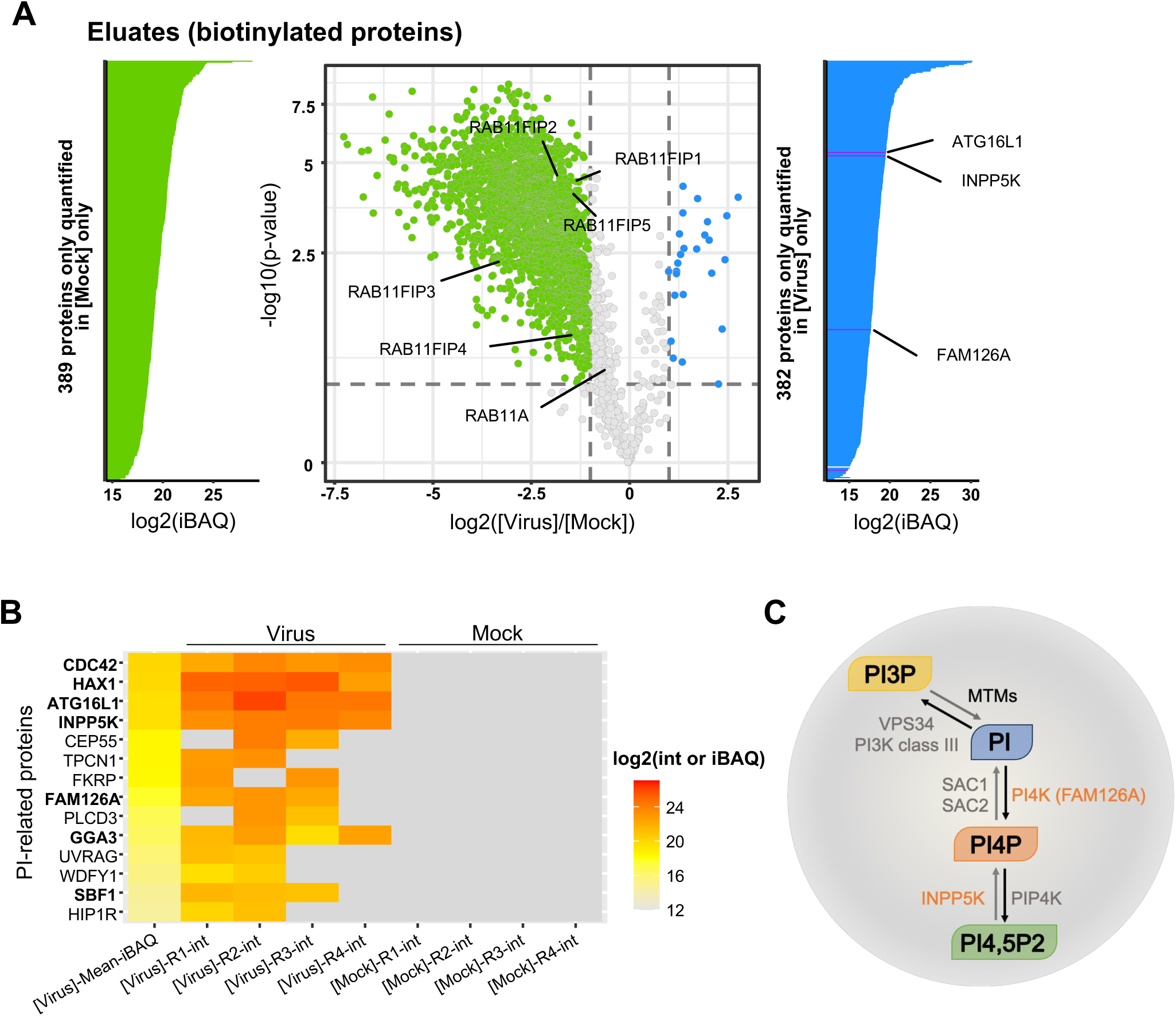
Viral-induced perturbations of RAB11A proximity interactome. **A.** Volcano plot showing the log2 fold change (x axis) and its significance (-log10(p-value, y axis) associated to a False Discovery Rate < 1%, for each protein (dots) in eluates from the RAB11A proximity labelling experiment. The log2 fold change refers to the enrichment in WSN-infected (n=4) versus mock-infected (n=4) samples. Blue and green dots represent proteins enriched in WSN-infected versus mock-infected samples, and proteins enriched in mock-infected versus WSN-infected samples, respectively. The iBAQ (intensity Based Absolute Quantification) plots are shown on the sides of the volcano plot for proteins that are present in WSN-infected samples only (blue) or in mock-infected samples only (green). **B.** Heat-map of cellular proteins associated with a GO term or a description containing the term “phosphatidylinositol”, according to the intensity values measured in the IAV-infected (n=4) or mock-infected (n=4) samples. On the left side of the heatmap, the “mean iBAQ in IAV” column represents the mean of the iBAQ values in the virus-infected samples. The iBAQ value approximates the abundance of a protein by dividing the (total precursor) intensities by the number of theoretically observable tryptic peptides of the protein. The log2 scale for intensity values/iBAQ values is shown on the right. **C.** Schematic representation of the enzymatic regulation of the PI3P/PI4P balance. Black and grey arrows represent the activities of the indicated kinases and phosphatases/phosphatase-related proteins, respectively. MTMs: myotubularins; INPP5K: inositol polyphosphate-5-phosphatase K; PI3K, PI4K, PI4PK: PI3-, PI4-and PI4P kinases, respectively; SAC1 and SAC2: PI4P phosphatases.

The A549-TurboID-RAB11A clonal cells were infected with the WSN virus at a MOI of 5 PFU/cell or mock-infected. At 9 hpi, biotin was added to the culture medium, cell lysates were prepared and biotinylated proteins were purified via streptavidin beads, as described in [28]. A fraction of the total lysates and the biotinylated eluates were then analysed by mass spectrometry-based proteomics.

Correlation analysis of the mass spectrometry data showed that the independent biological replicates were highly consistent, with Pearson correlation coefficients between the protein intensity values higher than 86.76% and 96.02% for the WSN- and mock-infected samples (n=4), respectively (**S3D Fig)**. Moreover, the distribution of protein intensity values across the different samples was very similar (**S3E Fig**). Gene Ontology (GO) enrichment analysis revealed a significant enrichment in GO terms related to membrane-associated proteins within the 3775 proteins found to be enriched in biotinylated eluates (**S4A Fig and S1 File**), which provides evidence for the specificity of our proximity labeling dataset.

Upon differential analysis of the biotinylated eluates, using a fold-change (FC) threshold of 2 and a false discovery rate (FDR) threshold of 1%, we found that 2382 and 407 cellular proteins were less abundant and more abundant, respectively, in the eluates derived from WSN-infected cells compared to the mock-infected controls (**Fig 3A**, green and blue color, respectively, and **S2 File**). Importantly, 1901 out of 2382 and 370 out of 407 proteins did not show the same trend of lower or greater abundance, respectively, when total lysates of WSN- and mock-infected cells were compared (**S4B Fig** and **S3 File**). Therefore, the variations in protein abundance observed in biotinylated eluates, whether they derive from infected or uninfected cells, most likely reflect specific, viral-induced changes. The large number of proteins with reduced or increased abundance demonstrate that the proximity interactome of RAB11A is strongly disrupted upon infection. Notably, several members of the RAB11-family interacting proteins known to be involved in the RAB11-mediated recycling pathway, i.e. RAB11FIP-1, -2, -3, -4 and -5, show reduced abundance (**Fig 3A**). This observation is fully in line with previous reports showing that the endosomal recycling function of RAB11 is impaired in IAV-infected cells [8,9].

Among the 370 proteins with increased abundance in RAB11A proximity interactome, we found a significant enrichment of GO terms related to mitochondrial proteins, notably mitochondrial ribosomal proteins (**S2 File**). This enrichment could possibly relate to a previous report that IAV infection alters the morphodynamics of mitochondria [29]. Interestingly, several proteins known to regulate the phosphoinositides (PIs) metabolism were exclusively quantified in biotinylated eluates derived from IAV-infected cells but not in those derived from mock-infected cells (**Fig 3B**). Seven proteins (CDC42, HAX1, ATG16L1, INPP5K, FAM1261 and SBF1) were notably identified in at least three out of the four replicates of infected biotinylated eluates*, i.e.* with a good reproducibility. Among these, two proteins are enzymatic regulators of the balance between the mono-phosphates PI3P and PI4P (**Fig 3C**), i.e. FAM126A (also known as HYCC1, a subunit of the phosphatidylinositol 4-kinase complex PI4KIIIA [30]) and INPP5K (a 5-P phosphatase producing mostly PI4P [31]), while ATG16L1, an autophagy-related protein, was shown to modulate the production and trafficking of PIP4 [32]. These findings suggest a potential link between PI4P homeostasis and RAB11A-mediated transport of vRNPs.

### IAV infection induces an increase in PI4P production at the ER

Our finding that upon IAV infection, RAB11A environment becomes enriched in proteins associated with PIs metabolism is consistent with the fact that PIs, especially mono-phosphates such as PI3P and PI4P, were reported to be closely associated with RAB11A-positive membranes [33] and to play an essential role in stress-dependent regulation of endomembrane morphodynamics [15,33,34]. These observations prompted us to investigate a possible alteration of the PIs balance upon IAV infection. We analyzed the global levels of PI3P and PI4P by immunofluorescence, and compared the mean PI3P or PI4P signal intensity per cell in WSN-infected compared to mock-infected A549 cells. At 8 hpi, using a FYVE-GST purified peptide [35] to label PI3P-positive membranes, we detected a sharp decrease of the PI3P signal in infected cells compared to control cells (p < 0.0001) (**Fig 4A-B**, left panels). In contrast, using an anti-PI4P antibody, we detected a stronger PI4P signal in infected cells compared to control cells (p < 0.0001) (**Fig 4A-B**, right panels). Then, the subcellular distribution of the PI4P signal with respect to ER membranes was monitored using U2OS-mEmerald-Sec61ß cells. In mock-infected cells, the few PI4P positive punctae are dispersed throughout the cytoplasm, whereas PI4P punctae formed upon IAV infection at 8 hpi are mostly associated with the ER (**Fig 4C-D**).

**Legend of Fig 4.**
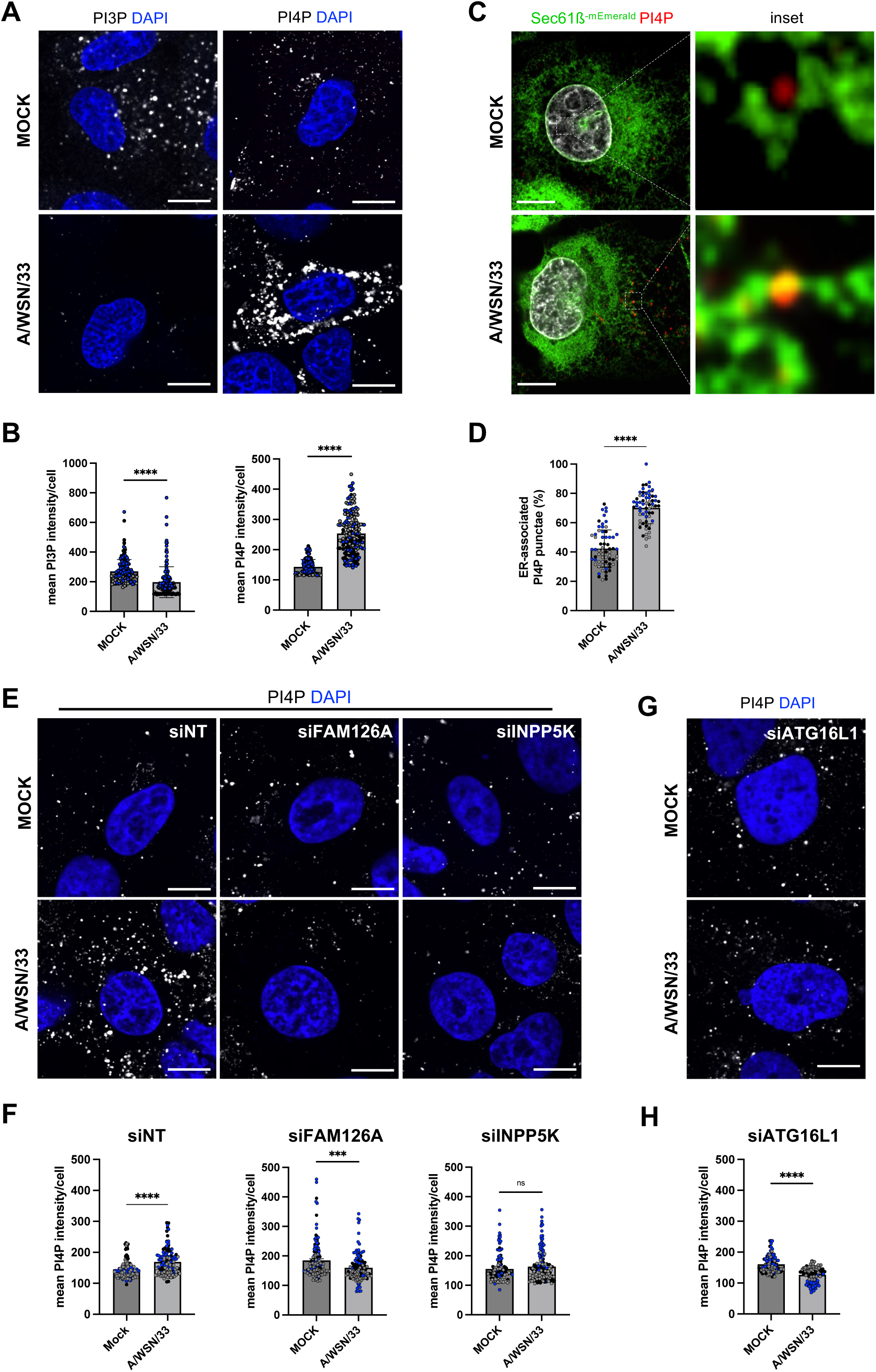
Viral-induced perturbations of the PI3P/PI4P balance. **A.** A549 cells were infected with WSN at a MOI of 5 PFU/cell for 8 h, or mock-infected. Fixed cells were stained for PI3P or PI4P. Nuclei were stained with DAPI and cells were imaged with a confocal microscope. Scale bar: 10 µm. **B.** A549 cells treated as in A were analyzed with the Fiji software to determine the mean intensity of the PI3P or PI4P signal per cell. Each dot represents one cell, and the data from three independent experiments are shown (black, grey and blue dots). The median and standard deviation values are represented (100-150 cells per condition). **\*\*\*\*** : p-value <0.0001, unpaired t-test. **C.** U2OS-Sec61ß-mEmerald cells were infected with WSN at a MOI of 5 PFU/cell for 8 h, or mock-infected. Fixed cells were stained for PI4P and nuclei were stained with DAPI. Cells were imaged with a confocal microscope. Scale bar: 5 µm. **D.** U2OS-Sec61ß-mEmerald cells treated as in C were analyzed with the Fiji software to determine the percentage of the total PI4P puncta associated to ER in individula cells. Each dot represents one cell, and the data from three independent experiments are shown (black, grey and blue dots). The mean and standard deviation values are represented as histograms (63-66 cells per condition). **\*\*\*\*** : p-value <0.0001, unpaired t-test. **E and G.** A549 cells were treated with control non-target (NT) siRNAs or with siRNAs targeting FAM126A, INPP5K (E) or ATG16L1 (G) for 48 h, and subsequently infected with WSN at a MOI of 5 PFU/cell for 8 h, or mock-infected. Fixed cells were stained for PI3P or PI4P. Nuclei were stained with DAPI and cells were imaged with a confocal microscope. Scale bar: 10 µm **F and H.** A549 cells treated as in E and G, respectively were analyzed with the Fiji software to determine the mean intensity of the PI3P or PI4P signal per cell. Each dot represents one cell, and the data from three independent experiments are shown (black, grey and blue dots). The median and standard deviation values are represented (100-150 cells per condition). **\*\*\*** : p-value <0.001, **\*\*\*\*** : p-value <0.0001, ns:non significant, unpaired t-test.

We then performed siRNA-mediated knock-down of the two RAB11A proximity-labeling hits known to enzymatically regulate the PIs metabolism in favor of PI4P synthesis, FAM126A and INPP5K (**S5B-C Fig**), and analyzed the global levels of PI4P upon IAV infection. Interestingly, the depletion of one or the other of these proteins abolishes the PI4P increase that we reported in IAV-infected cells (**Fig 4E-F**). Similar results were obtained by knocking-down the autophagic ATG16L1 protein, also detected in the vicinity of RAB11A by TurboID upon infection and known to be involved in PI4P turnover [32] (**Fig 4G-H** and **S5A,S5C Figs**).

In summary, our data show that IAV infection induces an increase in PI4P production at the ER, and this increase depends – at least partially – on FAM126A , INPP5K and ATG16L1.

### ATG16L1 and RAB11A control the abundance, proximity to vRNPs and/or turnover of PI4P pools in AIV-infected cells

As both PI4P (**Fig 4C-D**) and vRNPs (**Fig 2**) are detected at the vicinity of ER membranes in infected cells, we sought to investigate their physical and functional interplay. To this end, we examined the impact of ATG16L1 or RAB11A depletion on the production of PI4P and the distribution of PI4P relative to ER membranes and vRNPs, in IAV-infected U2OS-mEmerald-Sec61ß cells (**Fig 5A-D**). The depletion of ATG16L1 led to a significant reduction of the number of PI4P punctae per cell (**Fig 5B**), in agreement with our previous observations in IAV-infected A549 cells (**Fig 4G-H**), with a conserved percentage of ER-associated PI4P punctae (**Fig 5C**). The depletion of RAB11A also led to a significant reduction of the number of PI4P punctae per cell (**Fig 5B**), but unlike the depletion of ATG16L1, it resulted in a moderate but significant increase in the percentage of PI4P punctae associated with the ER in infected cells (86 + 11 %, compared to 70+ 11 % in control cells, p < 0.0001) (**Fig 5C**). These observations suggests that i) both ATG16L1 and RAB11A contribute to the amplification of PI4P pools at the vicinity of the ER, and ii) RAB11A contributes to the turnover of these pools, possibly in relation with the biogenesis of RAB11A-positive, vRNP-coated vesicles from PI4P-enriched ER subdomains. To further assess this hypothesis, we measured the percentage of PI4P punctae associated with NP-positive punctae. As shown in **Fig 5D**, it was 55 + 14 % in control IAV-infected cells, and was reduced to 20 + 15 % in ATG16L1-depleted cells (p<0.0001). This parameter could not be measured in RAB11A-depleted cells because of the diffuse pattern of the NP signal. Finally, we analyzed the distribution of ATG16L1, and observed that it relocalizes from a predominantly cytoplasmic distribution in mock-infected cells (**Fig 5E**, upper panels) to ER membranes in WSN-infected cells (**Fig 5E**, lower panels). Altogether our data suggest that ATG16L1 not only contributes to the amplification of PI4P pools at the vicinity of the ER, but also controls their proximity with vRNPs.

**Legend of Fig 5.**
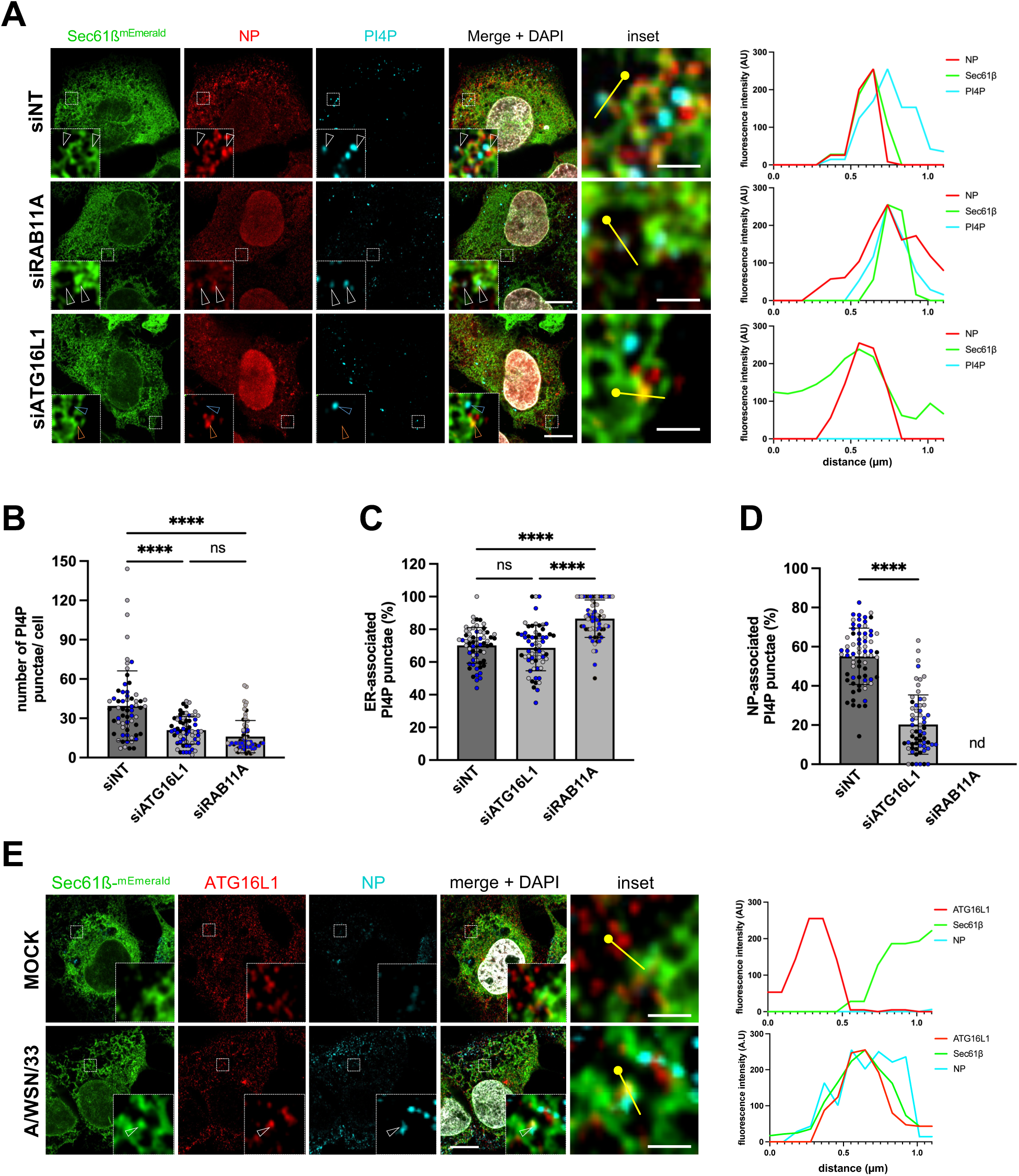
Impact of RAB11A or ATG16L1 depletion on viral-induced production and localization of PI4P. **A.** U2OS-Sec61ß-mEmerald cells were treated with control Non-Target (NT) siRNAs or with siRNAs targeting RAB11A or ATG16L1 for 48 h, and subsequently infected with WSN at a MOI of 5 PFU/cell for 8 h. Fixed cells were stained for PI4P and NP, and nuclei were stained with DAPI. Cells were imaged with a confocal microscope. Whole cell scale bar: 5 µm; insets scale bar: 1 µm; arrowheads: NP-Sec61ß-PI4P co-distribution. For each condition, the graph on the right corresponds to a fluorescence intensity profile for NP (red), Sec61ß (green) and PI4P (cyan) along the yellow line drawn in the inset, starting from the knob. **B-D.** U2OS-Sec61ß-mEmerald cells treated as in (A) were analyzed with the Fiji software to determine the number of PI4P puncta (B), the percentage of PI4P puncta associated to the ER (C) and the percentage of PI4P puncta associated to NP puncta (D) in individual cells. Each dot represents one cell, and the data from three independent experiments are shown (black, grey and blue dots). The mean and standard deviation values are represented as histograms (62-70 cells per condition). (B-C) **\*\*\*\*** : p-value <0.0001, ns: not significant, two-way ANOVA. (D) **\*\*\*\*** : p-value <0.0001, unpaired t-test. nd: not done, because the diffuse cytoplasmic distribution of NP signal in cells treated with the siRNA targeting RAB11A precludes analysis of the percentage of PI4P puncta associated to NP puncta. **E.** U2OS-Sec61ß-mEmerald cells were infected with WSN at a MOI of 5 PFU/cell for 8 h, or mock-infected. Fixed cells were stained for ATG16L1 and NP, and nuclei were stained with DAPI. Cells were imaged with a confocal microscope. Whole cell scale bar: 5 µm; insets scale bar: 1 µm. arrowheads: NP-Sec61ß-PI4P co-distribution. For each condition, the graph on the right corresponds to a fluorescence intensity profile for Sec61ß (green), ATG16L1 (red) and NP (cyan) along the yellow line drawn in the inset, starting from the knob.

### ATG16L1-depletion or drug-mediated inhibition of PI4KIIIA activity leads to reduced viral progeny release

Upon infection with the WSN virus at a low MOI (0.001 PFU/cell), ATG16L1-depleted cells show a ∼one-log reduction of the production of infectious particles compared to control cells, while RAB11A-depleted cells show a ∼two-log reduction (**Fig 6A**). Upon infection at a high MOI (5 PFU/cell) and immunostaining for the NP protein at 8 hpi, the percentage of cells showing an accumulation of NP signal at the vicinity of the plasma membrane is reduced from ∼34% to 13 % (p < 0.05) (**Fig 6B-C**), suggesting that vRNP transport to the plasma membrane is delayed. These observations indicate that ATG16L1 is involved at late stages of IAV life cycle and promotes vRNP trafficking and the production of infectious viral particles. Taken together with our previous data, this proviral activity of ATG16L1 is most likely mediated by the control of local PI4P production on ER membranes.

**Legend of Fig 6.**
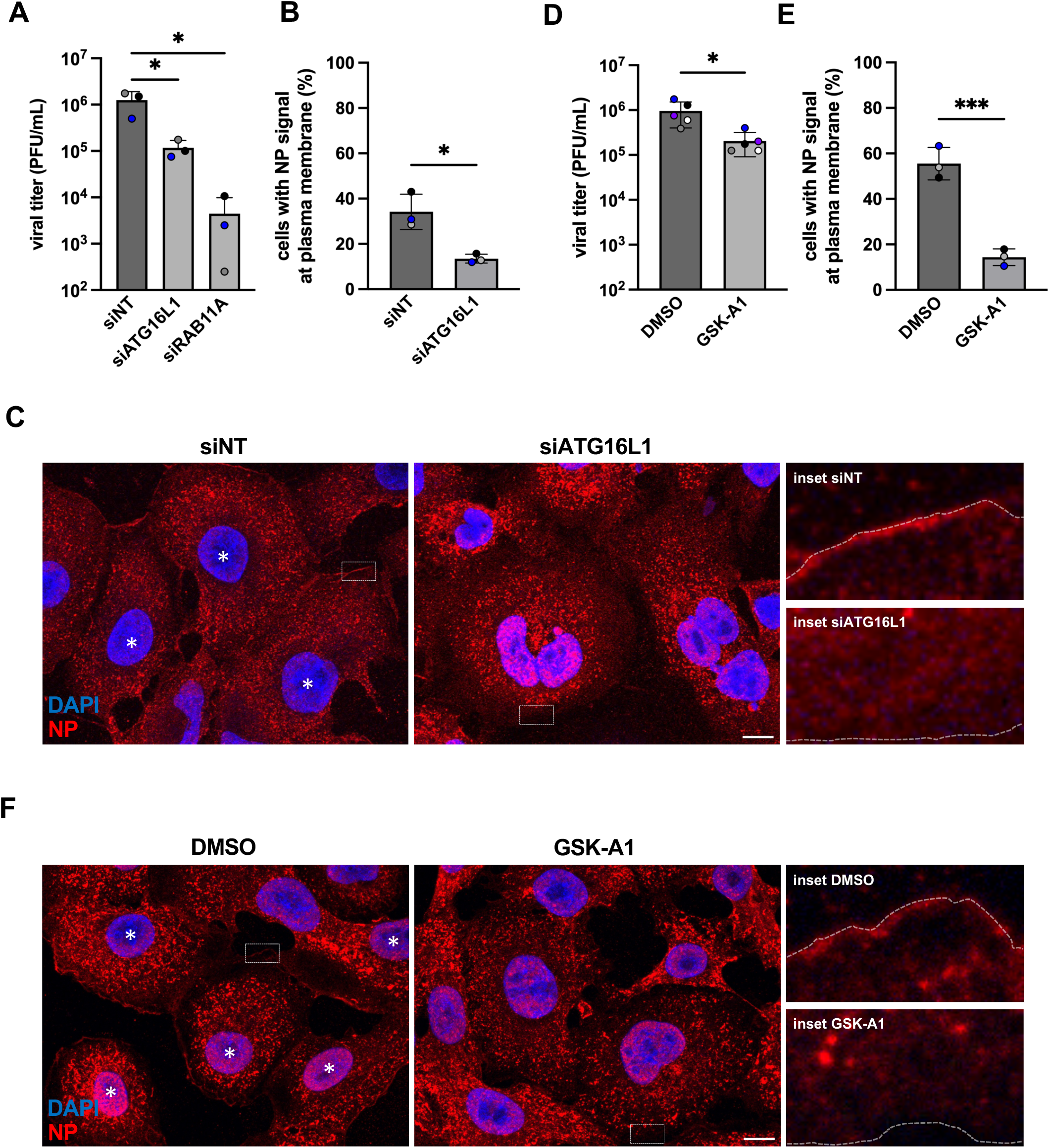
Impact of ATG16L1 depletion or GSK-A1 treatment on vRNP egress and viral progeny. **A.** A549 cells were treated with the indicated siRNAs for 48 hours and subsequently infected with WSN at a MOI of 0.01 PFU/mL. At 24 hpi, the supernatants were collected and the infectious titer were determined by a plaque assay. The mean +/-SD of three independent experiments is shown. **\*** : p-value <0.05, paired t-test. **B-C.** A549 cells were treated with the indicated siRNAs for 48 hours and subsequently infected with WSN at a MOI of 5 PFU/mL. At 8 hpi, fixed cells were stained for NP and nuclei were stained with DAPI. Cells were imaged with a confocal microscope. (B). The percentage of cells showing an accumulation of NP signal at the plasma membrane is represented (178-288 cells per condition) **\*** : p-value <0.05, paired t-test. (C) Representative cells are shown, and are labeled with a white star when an accumulation of NP signal at the plasma membrane is detected. Whole cell scale bar: 10 µm; insets scale bar: 1 µm. The dotted white line in the insets delineate the cell border. **D.** A549 cells were infected with WSN at a MOI of 5 PFU/mL. Two hours later, the GSK-A1 drug was added at a final concentration of 100 nM. DMSO at the same concentration was used as a control. At 6 hpi, the supernatants were collected and the infectious titer were determined by a plaque assay. The mean +/- SD of 5 independent experiments is shown. **\*** : p-value <0.05, paired t-test. **E-F.** A549 cells were treated as in D. At 6 hpi, fixed cells were stained for NP and nuclei were stained with DAPI. Cells were imaged with a confocal microscope. (E). The percentage of cells showing an accumulation of NP signal at the plasma membrane is represented (166-208 cells per condition) **\*\*\*** : p-value <0.001, paired t-test. (F) Representative cells are shown and labeled as in (C). Whole cell scale bar: 10 µm; insets scale bar: 1 µm. The dotted white line in the insets delineate the cell border.

To further confirm the importance of PI4P for the late stages of the viral cycle, we examined the impact of GSK-A1-mediated depletion of PI4P. GSK-A1, a selective inhibitor of the PI4KIIIA kinase complex which includes FAM126A [36], was added 2 hours post-infection with the WSN virus at a high MOI. At 6 hpi, the amount of infectious viral particles produced in the supernatant is reduced ∼4.5 -fold (p<0.05) in GSK-A1-treated cells compared to DMSO-treated cells (**Fig 6D**), whereas the levels of accumulation of viral proteins are unchanged (**S5D Fig**).The percentage of cells showing an accumulation of NP signal at the plasma membrane is reduced from ∼55% to 14% (p<0.001) (**Fig 6E-F**), similar to what observed in ATG16L1-depleted cells (**Fig 6B-C**). These observations further support the functional importance of the rewiring of intracellular PI4P to ER membranes for vRNP transport.

## Discussion

The first evidence that RAB11A is essential for the cytoplasmic transport of influenza vRNPs and the production of infectious virions were provided in the early 2010s [6,22–24]. Despite progress in characterizing these late stages of the viral life cycle (e.g. [9–11]), the precise mechanisms by which RAB11A is involved remain elusive. Here, we show that IAV infection strongly modifies the proximity interactome of RAB11A, contributes to an alteration of the balance between phosphoinosides PI3P and PI4P, and induces a cross-talk between RAB11A, ATG16L1 and PI4P at the vicinity of the ER, which promotes vRNP transport.

Among the > 1900 proteins whose abundance in RAB11A proximity interactome was decreased upon infection, we found the RAB11FIP-1, -2, -3, -4, and -5 proteins, i.e. RAB11A effector proteins essential to the endosomal recycling process. This finding is consistent with previous reports that the efficiency of RAB11A-dependent recycling pathway is altered in IAV-infected cells [8,9] and that depletion of RAB11FIP-2 or RAB11FIP-3 has no detectable impact on the production of infectious viral particles [23]. It reinforces the notion that in IAV-infected cells, the intracellular localization, topology and function of RAB11A-positive membranes are distinct from those of recycling endosomes [11].

In contrast, RAB11A proximity interactome in IAV-infected cells is enriched in proteins (FAM126A, INPP5K, ATG16L1) known to stimulate the production of PI4P, a key player of the stress-dependent regulation of endomembrane morphodynamics [33,34,37]. We show that concommitantly, IAV infection induces a significant decrease in PI3P as well as a significant increase of PI4P cellular levels, in a RAB11A-dependent and ATG16L1-dependent manner. Treatment with a selective inhibitor of the type III PI4KA kinase, which is regulated by FAM126A, results in a delayed accumulation of vRNPs at the plasma membrane and a reduction of the production of infectious viral particles, demonstrating the functional importance of PI4P for vRNP transport. Notably, the PI4P fraction associated with ER membranes represents ∼70% of the total PI4P pool in IAV-infected cells, compared to ∼40% in mock-infected cells. These very novel observations, taken together with the fact that RAB11A, ATG16L1 and vRNPs are localized at the vicinity of ER membranes upon infection (this study and [9–11]), point to the ER as an essential transport platform for vRNPs.

We confirm our previous observations that IAV infection induces the remodeling and tubulation of ER membranes all throughout the cell, in a RAB11A-independent manner [11]. The viral-induced depletion in PI3P, a phosphoinositide enriched in the membrane of early endosomes [38], might contribute to ER remodeling. Indeed, the reduction in PI3P levels induced by nutrient deprivation, a distinct type of cellular stress, was shown to result in a loss of contacts between early endosomes and the ER and a subsequent reshaping of the ER [39]. However, unlike the conversion of tubular membranes to sheets induced by nutrient deprivation [39], we observed an extension and tubulation of sheet membranes towards the periphery of IAV-infected cells. ER stress signalling, which was shown to be induced by the expression of IAV glycoproteins, and balanced by the NS1 viral protein through its host protein shutoff activity [40,41], could possibly contribute to ER remodeling. In addition, or alternatively, the hemagglutinin and neuraminidase could directly trigger ER remodeling by inducing membrane zippering, as recently shown by *in situ* cryo-electron tomography [12].

Based on super-resolution microscopy and statistics-based quantification, we provide evidence that vRNPs tend to accumulate at the vicinity of ER membranes in RAB11A-depleted cells. Uncovering the RAB11A-independent mechanism by which vRNPs are targeted to the ER is beyond the scope of this study. Notably, we show that ∼ 60% of the PI4P punctae present at the ER membranes upon IAV infection co-distribute with vRNPs. Depletion of RAB11A results in a moderate but significant increase of the PI4P associated with ER membranes, suggesting the existence of a RAB11A-PI4P cross-talk coupled with PI4P turnover. There is multiple evidence that cross-talks between phosphoinositides and RAB GTPases can control membrane remodeling and trafficking [16]. In particular, PI4P was shown to recruit the RAB11A effector KIF13A and the membrane-shaping factor BLOC1 to induce the remodeling of RAB11A-positive sorting endosomes into recycling endosomal tubules [42]. KIF13A and the BLOC1 subunit 1 (BLOC1S1) are detected at the proximity of RAB11A in mock-infected cells; they are significantly less abundant (KIF13A) or undetected (BLOC1S1) at the proximity of RAB11A in IAV-infected cells (**S3 File**), which reinforces the notion that the environment of RAB11A membranes is strongly altered upon infection. We speculate that in the context of IAV-infected cells, a RAB11A-PI4P cross-talk at the vicinity of ER subdomains induces the recruitment of distinct RAB11A effectors and/or membrane-shaping factors, and leads to the biogenesis of RAB11A-positive, vRNP-coated transport vesicles. This hypothesis is consistent with the observation that vRNPs and RAB11A-positive membranes concentrate in liquid condensates at the vicinity of the ER [9,10]. Contact sites between ER membranes and RAB11A-positive membranes, or possibly membrane fusion events, are likely involved and mediate the transfer of vRNPs from the vicinity of the ER membrane to the transport vesicles.

Our data provide evidence for a pivotal role of ATG16L1 in coordinating RAB11A, vRNPs and PI4P-enriched membranes in the process of vRNP transport. Indeed, upon infection of ATG16L1-depleted cells, the cellular PI4P levels, the proportion of PI4P punctae associated with vRNPs, the proportion of cells showing an accumulation of vRNPs at the plasma membrane at 8 hpi, and the production of infectious viral particles, were all significantly reduced compared to control ATG16L1-expressing cells. Beyond its role in autophagy, ATG16L1 appears as an a important player in the regulation of the endomembrane system in response to stress [43]. It was notably shown to be involved in the biogenesis of the primary cilium, a sensor of external chemical and mechanical stress, by targeting the INPP5E phosphatase at the cilium membrane to produce PI4P [32]. Our data indicate that in the context of IAV-infected cells, ATG16L1 controls the identity of ER membrane subdomains in terms of PI4P content and PI4P proximity with vRNPs, and cooperates with RAB11A present at the vicinity of the ER. Although a cooperation between ATG16L1 and RAB11A-positive membranes has been described in the context of the canonical autophagic response to starvation stress [44], several observations suggest that in the context of an IAV infection, the mode of action of ATG16L1 could be unrelated to canonical autophagy. Indeed, IAV infection was shown to initiate the autophagy process but to prevent its full completion [45]. Moreover, important partners of ATG16L1 in canonical authophagy, such as ATG5-ATG12 [34], and WIPI2, shown to be recruited by RAB11A for ATG16L1-mediated assembly of autophagosomes [44] were not detected in the RAB11A interactome along with ATG16L1. Finally, another component of the autophagy machinery, ATG9A, was recently shown to facilitate the formation of liquid condensates enriched in vRNPs and RAB11A-positive membranes independently from its function in autophagy, by regulating membrane-microtubule contacts [9].

Therefore, we propose that upon IAV infection, ATG16L1 acts as a stress sensor that coordinates with RAB11A to regulate the identity of endomembranes to ensure delivery of vRNP-coated membranes to the cell surface (**Fig 7**). The precise sequence of protein recruitment engaged in this process remains to be determined. Our findings extend to IAVs the notion that viruses have evolved strategies to modulate the metabolism and localization of cellular lipids. They also highlight the role of proteins of the autophagy machinery in regulating intracellular trafficking, and the highjacking of this function by pathogens.

**Legend of Fig 7.**
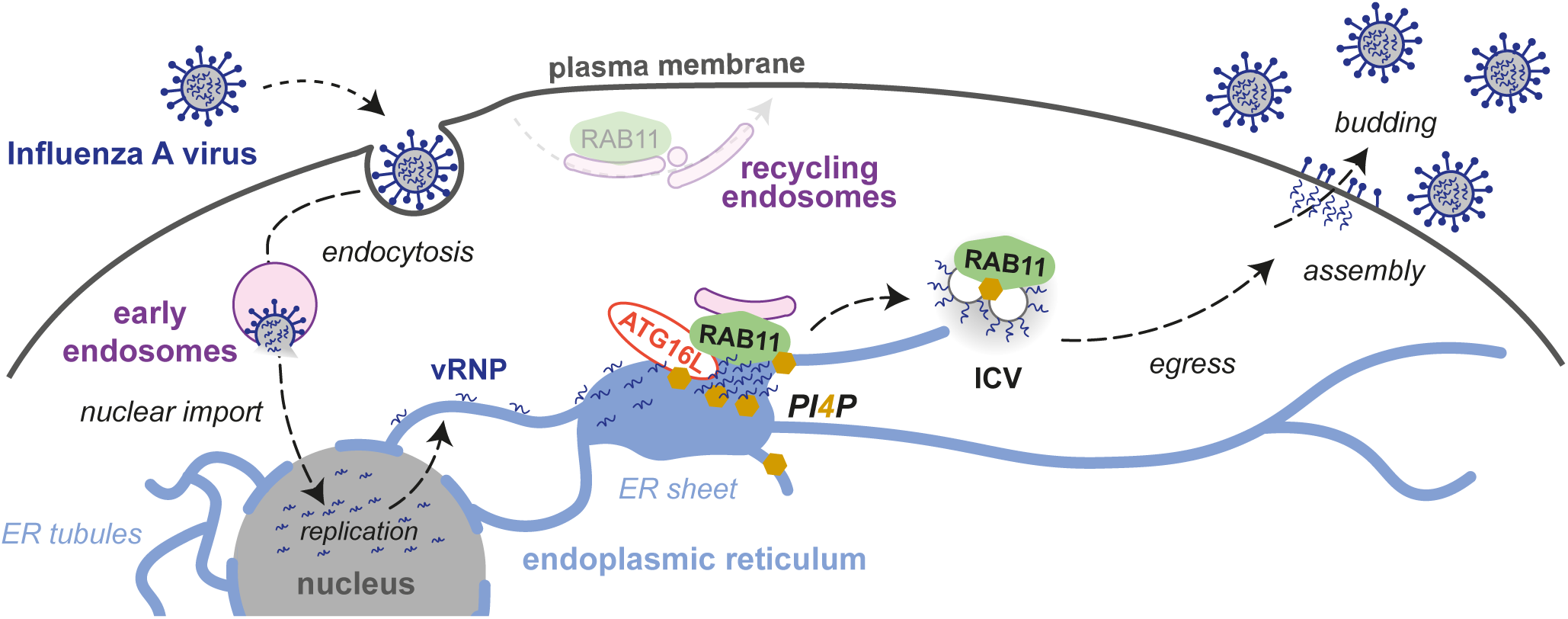
Model for influenza vRNP transport from the nucleus to the plasma membrane. Upon influenza virus entry, viral ribonucleoproteins (vRNPs) are imported into the nucleus and they serve as a template for the transcription and replication of the viral genome. The neosynthesized vRNPs are exported from the nucleus and get associated with the endoplasmic reticulum (ER) membrane. Viral infection promotes the remodeling of ER sheets and their extension towards the cell periphery. The recruitment of ATG16L1 and RAB11 to vRNP-enriched ER sites induces the local production of PI4P and favors the biogenesis of vRNP transport vesicles previously described as irregularly coated vesicles (ICVs) [9]. RAB11A mediates the transport of ICV egress towards the plasma membrane, where viral particles are assembled and bud from the cell surface.

## Materials and Methods

### Cells and viruses

HEK-293T (ATCC CRL-3216), A549 (kindly provided by Pr. M. Schwemmle, University of Freiburg) and U20S-Sec61ß-mEmerald cells (engineered as in [26], kindly provided by O. Schwartz, Institut Pasteur, Paris, France) were grown in complete Dulbecco’s modified Eagle’s medium (DMEM, Gibco) supplemented with 10% (v:v) fetal calf serum (FCS), 100 U/mL penicillin and 100 µg/mL streptomycin. Madin-Darby Canine Kidney (MDCK) cells, kindly provided by the National Reference Center for Respiratory Viruses (Institut Pasteur, Paris, France) were grown in Modified Eagle’s Medium (MEM) supplemented with 5% FCS, 100 U/mL penicillin and 100 µg/mL streptomycin. The recombinant viruses A/WSN/33 (WSN, [46]), WSN-PB2-Strep [47]), and A/Victoria/3/75 (VIC, [48]) were produced by reverse genetics as described in [47]. The Zika strain PF13 (kindly provided by V. M. Cao-Lormeau and D. Musso, Institut Louis Malardé, Tahiti Island, French Polynesia) was isolated from a viremic patient in French Polynesia in 2013. When indicated, cells were treated with the GSK-A1 drug (Sigma-Aldrich, 100nM) dissolved in dimethylsulfoxyde (DMSO, Sigma-Aldrich) from 2 to 6 hours post-infection.

### Plasmids and lentiviral transduction

Reverse genetics plasmids for the WSN virus were kindly provided by G. Bronwlee (Oxford University, UK). Reverse genetics plasmids for the VIC virus [48] were developed in J. Ortin’s group and kindly provided by W. Barclay (Imperial College London, UK). To generate the lentiviral pLVX-3xHA-TurboID-RAB11A plasmid, the 3xHA-TurboID coding sequence was amplified by PCR using the plasmid 3xHA-TurboID-NLS-pCDNA3 (a gift from A.Ting, Standford University, USA, Addgene #107171 [27] as a template, and subcloned instead of mCherry into pLV-CMV-mCherry-RAB11A-IRES-puro, an unpublished derivative of the pLV-CMV-eGFP lentiviral vector (Duke vector core, Duke University). To produce the corresponding lentiviruses, 10^7^ HEK-293T cells were plated in 10 cm dishes, before co-transfection the next day with 15 µg of the pLVX-3xHA-TurboID-RAB11A plasmid along with 10 and 5 µg of the pΔ8.74 (Addgene #22036) and pMD2.G (Addgene #12259) packaging plasmids, respectively, using polyethylenimine (PEI, Polysciences, Inc., 3 µL of PEI at 1 mg/mL for 1 µg of DNA). At 72 hpt the conditionned media was passed through a 0.45μm-pore size filter and placed over A549 cells. At 48 h post-transduction, cells were split and cultured in puromycin (1 µg/mL) containing media to select for a polyclonal subpopulation of transduced cells. Cellular clones were isolated upon limiting dilution in 96-well plates. The resulting polyclonal and clonal cells were assessed for expression of 3xHA-TurboID-RAB11A by western blotting.

### Affinity purification of biotinylated proteins

The protocol for TurboID-mediated proximity labeling was an adaptation from the protocol described in [28]. Briefly, A549-TurboID-RAB11A cells were seeded in 75 cm^2^ dishes (10^7^ cells/dish) and infected with the WSN virus at a MOI of 5 PFU/cell or mock-infected. At 8 hpi, the culture medium was replaced with 10 mL of prewarmed (37°C) medium supplemented with 50 µM biotin (Sigma Aldrich, B4501). The cells were incubated at 37°C for 10 min, washed 4 times with 2 mL ice-cold PBS, and dry-frozen at -80°C for at least 1h 30 min. They were then lysed in 1.2 mL of RIPA buffer, incubated for 10 min on ice, and centrifuged for 10 min at 13,000g at 4°C. The supernatants were collected, an aliquot of 120 µL was frozen at -80°C and the remaining was incubated with 125 µL of magnetic streptavidin beads (ThermoFisher #88817, pre-washed according to the supplier’s recommendations) overnight at 4°C on a rotating wheel. The beads were washed at room temperature using a magnet, twice with 1 mL of RIPA buffer (2 min), once with 1 mL of KCl 1M (2 min), once with 1mL Na_2_CO_3_ (10 s), once with 1mL urea in 10mM Tris HCl (10 s), and twice with 1mL of distilled water (2 min). Upon centrifugation, 950 µL of water was removed, and the beads covered with the remaining 50 µL were kept frozen at -80°C until tryptic digestion on beads and LC-MS/MS analysis.

### LC-MS/MS sample preparation and data acquisition

#### Whole cell proteome

Proteins from cell lysates were reduced using 5 mM dithiothreitol (DTT, Sigma-Aldrich #43815) for 30 min at 25°C with agitation. Following reduction, the proteins were alkylated with 20 mM iodoacetamide (IAA, Sigma-Aldrich #I114) for 30 min at 25°C under agitation. Protein isolation and digestion were carried out using the Single-Pot Solid-Phase-enhanced Sample Preparation (SP3) method, with minor modifications from the original protocol as described by Hughes et al. [49]. In brief, SP3 beads were prepared by mixing hydrophilic and hydrophobic Sera-Mag SpeedBeads (GE Healthcare, Chicago, IL, USA) in a 1:1 (v/v) ratio, washed three times with water, and reconstituted to a final concentration of 50 µg/µL. To the protein samples, 8 µL of prepared beads were added, followed by the addition of anhydrous acetonitrile (ACN) to a final concentration of 75% (v/v). The samples were agitated in a thermomixer at 800 rpm for 30 min at room temperature. After 1 min on a magnet, the supernatant was removed, and the beads were washed twice with 80% ACN and once with 100% ACN. For digestion, 100 mM ammonium bicarbonate (ABC, pH 8.0) containing sequencing-grade modified trypsin (Promega, #V5111) was added to the bead-bound proteins at a protein-to-enzyme ratio of 30:1. The samples were incubated for 12 h at 37°C. After digestion, the supernatant containing peptides was collected into a new tube following 1 min on a magnet. The beads were then washed with water, mixed for 10 min at room temperature, and the supernatant was pooled with the initial peptide solution. The resulting peptides were acidified with formic acid (FA) to a final concentration of 1%.

#### Eluates enriched in biotinylated proteins

The processing of interactomics samples was performed as per the protocol detailed by Cho et al. (2020) [28]. Briefly, protein attached to the beads were incubated with a digestion buffer consisting of 1 M urea, 1 mM DTT, and 50 mM Tris-HCl (pH 8.0) containing 0.4 µg trypsin, for 1 h at 25°C with agitation. The supernatants were collected into fresh tubes, and the beads were washed twice with a mild denaturing buffer containing 2 M urea and 100 mM ABC. The supernatants were pooled, reduced with 5 mM DTT for 30 min at 25°C under agitation, and alkylated with 20 mM IAA for 30 min at 25°C with agitation. The samples were diluted 2-fold with 100 mM ABC, and an additional 0.5 µg of trypsin was added for further digestion over 16 h at 37°C with agitation. The resulting peptides were acidified to 1% with formic acid (FA).

#### Desalting

Digested peptides were desalted using C18 cartridges (Agilent Technologies, 5 μL bead volume, 5190-6532) and eluted with ACN 80 %, FA 0.1 %. Finally, the peptide solutions were speed-vac dried and resuspended in ACN 2%, FA 0.1% buffer. Only for global proteome samples, absorbance at 280 nm was measured with a NanodropTM 2000 spectrophotometer (Thermo Scientific) to inject an equivalent of DO = 1.

#### Mass Spectrometry

A nanochromatographic system (Proxeon EASY-nLC 1200 - Thermo Fisher Scientific) was coupled on-line to a Q Exactive^TM^ Plus Mass Spectrometer (Thermo Fisher Scientific) using an integrated column oven (PRSO-V1 - Sonation GmbH, Biberach, germany). For each sample, peptides were loaded into a capillary column picotip silica emitter tip (home-made column, 40cm x 75 µm ID, 1.9 μm particles, 100 Å pore size, Reprosil-Pur Basic C18-HD resin, Dr. Maisch GmbH) after an equilibration step in 100 % buffer A (H_2_O, FA 0.1%). Peptides from global proteome samples were injected at constant quantity and peptides from enriched biotinylated proteins at constant volume. Peptides were eluted with a multi-step gradient from 5 to 25 % buffer B (80 % ACN, FA 0.1 %) during 95 min, 25 to 40 % during 15 min and 40 to 95 % during 10 min at a flow rate of 300 nL/min over 130 min. Column temperature was set to 60°C.

MS data were acquired using Xcalibur software using a data-dependent Top 10 method (Global proteome) or a Top 5 method (Biotin enriched proteins) with survey scans (300-1700 m/z) at a resolution of 70,000 and MS/MS scans (fixed first mass 100 m/z) at a resolution of 17,500. The AGC target and maximum injection time for the survey scans and the MS/MS scans were set to 3.0E6, 20 ms and 1E6, 60 ms (global proteome) or 100 ms (enriched biotinylated proteins) respectively. The isolation window was set to 1.6 m/z and normalized collision energy fixed to 28 for HCD fragmentation. We used a minimum AGC target of 1.0E4 for an intensity threshold of 1.7E5 (global proteome) or 1.0E5 (enriched biotinylated proteins). Unassigned precursor ion charge states as well as 1, 7, 8 and >8 charged states were rejected and peptide match was disabled. Exclude isotopes was enabled and selected ions were dynamically excluded for 45 sec (global proteome) or 30 sec (enriched biotinylated proteins).

#### Protein Identification and Quantification

MS Raw files were uploaded into MaxQuant software (version 2.0.3.0), where A549 data were searched against the Uniprot *homo sapiens* database (20,360 proteins the 13/12/2021), and WSN virus data were searched against the Influenza A virus database (10 proteins the 27/07/2022), both of which were modified to contain target sequences for TurboID. Methionine oxidation, protein N-terminal acetylation, and lysine biotinylation were variable modifications; cysteine carbamidomethylation was a fixed modification; the maximum number of modifications to a protein was 5. The minimum peptide length was set to 7 amino acids, with a maximum peptide mass of 8000 Da. Search was performed with trypsin as specific enzyme with a maximum number of two missed cleavages. Identifications were matched between runs within replicates of a same condition. A false discovery rate (FDR) cutoff of 1 % was applied at the peptide and protein levels. Peptides were quantified using unique and razor peptides. The mass spectrometry proteomics data have been deposited to the ProteomeXchange Consortium via the PRIDE partner repository [50] with the dataset identifier PXD053858.

#### Statistical analysis of mass spectrometry data

For both the global proteome and enriched biotinylated protein samples, Maxquant intensities were uploaded into our R package. To find the proteins more abundant in one condition than in another, the intensities quantified using Maxquant were compared. Reverse hits, potential contaminants, and proteins not well identified with a 1% FDR (“Only identified by site”) were first removed from the analysis. Only proteins identified with at least one peptide that is not common to other proteins in the FASTA file used for the identification (at least one “unique” peptide) were kept. Additionally, only proteins quantified in at least two replicates of one of the two compared conditions were kept for further statistics. Proteins without any value in one or the other biological condition have been considered as proteins quantitatively present in a condition and absent in another. They have therefore been set aside and considered as differentially abundant proteins. They have been displayed in barplots and ranked from the one with the highest IBAQ (intensity-based absolute quantification) value to the lowest on the side of volcano plots (**Fig 3A** and **S4 Fig**). An IBAQ value is a measure of protein abundance [51]. After this step, intensities of the remaining proteins (quantified in both conditions) were first log-transformed (log2). Next, intensity values were normalized by median centering within conditions (section 3.5 in [52]). Missing values were imputed using the impute.slsa function of the R package imp4p [53]. Statistical testing was conducted using a limma t-test thanks to the R package limma [54]. An adaptive Benjamini-Hochberg procedure was applied on the resulting p-values thanks to the function adjust.p of the cp4p R package [55] using the robust method described in [56] to estimate the proportion of true null hypotheses among the set of statistical tests. These proteins were displayed in volcano plots (**Fig 3A** and **S4 Fig**). The final set of proteins of interest is composed of the proteins which are differentially expressed according to this statistical analysis, and those which are absent from one condition and present in another (**S2 and S3 Files**).

Functional enrichment analyses of terms and pathways in the proteins of interest have been determined using the app stringApp [57] of Cytoscape [58]. Different backgrounds have been used for the enrichment tests: either the list of proteins identified in all the replicates of biotinylated eluates or the list of proteins identified in all the replicates of biotinylated eluates and total lysates. A significantly low p-value means the proportion of proteins related to a term or pathway is significantly superior in the considered list than in the used background. The PIP-related proteins displayed in the heatmap of **Fig 3B** were selected because their description or GO terms mention the keyword “phosphatidylinositol”.

#### Immunostaining for proteins

7.5×10^4^ A549- or U2OS-derived cells were seeded on glass coverslips (13mm diameter, #1.5, Epredia, CB00130RAC20MNZ0) in 24-well plates, and infected 24 h later with the WSN of VIC virus at a MOI of 5 PFU/cell. At the indicated time-points, cells were fixed with PBS-4% PFA (Fisher scientific, 47377) for 15 min and washed 3 times with 1mL PBS. For most protein immuno-stainings, cells were incubated in 50mM NH_4_Cl (Sigma Aldrich)-PBS for 10 min at room temperature to quench free aldehyde groups from PFA. Cells were washed 3 times in 1mL PBS and incubated in a Triton X100-containing permeabilization and blocking solution (PBS supplemented with 0.1% Triton X-100, 5% Normal Donkey serum) for 1h at room temperature. Cells were then incubated overnight at 4°C with primary antibodies directed against RAB11 (Invitrogen 71-5300, 1:100), CLIMP63 (R&D systems AF7355, 1:500), ATG16L1 (MBL PM040, 1:200), the HA tag (Invitrogen 26183, 1:100), or the influenza NP (BioRad MCA 400 or Invitrogen PA5-32242, 1:1000). Cells were then washed 3 times in 0,05% Tween 20-PBS and stained with an appropriate secondary antibody conjugated to an Alexa Fluor (AF) dye (Jackson Immunoresearch 715-545-150; 711-545-152 (AF488); 715-165-150 (Cy3); 711-606-152 (AF647), ThermoFisher A-21436 (AF555) 1:400) and with DAPI (ThermoFisher, 62248, 1 µg/mL) diluted in the same permeabilization and blocking solution. When indicated, DY-488-conjugated Strep-Tactin XT (IBA Lifesciences 2-1562-050, 1:200) was added in the same solution. Cells were then washed 3 times in 0,05% Tween 20-PBS, 1 time with 1mL PBS and 1 time with water. Coverslips were mounted on glass slides with Fluoromount-G antifade reagent (Invitrogen, 00-4958-02).

In the case of dual stainings of ER markers CLIMP63 and RNT3 (Santa Cruz sc-374599, 1:150), together with the influenza NP or Zika NS3 antigen (anti-NS3 antibody [59] kindly provided by Andres Merits, 1:1000), cells were incubated in a saponin-containing permeabilization and blocking solution (PBS supplemented with 0.1% saponin, 1% Bovine Serum Albumin) for 30 min at room temperature. Cells were washed 3 times in 1mL PBS for 5 min at room temperature. Free aldehyde groups from PFA were quenched using the same permeabilization and blocking solution supplemented with 20 mM glycine (Sigma Aldrich) for 15 min at room temperature. Cells were then incubated overnight at 4°C with primary antibodies diluted in permeabilization and blocking solution. Cells were washed 3 times in 1 mL PBS for 5 min at room temperature prior to incubation with appropriate AF-conjugated secondary antibodies and DAPI diluted in permeabilization and blocking solution for 1h at room temperature. Cells were washed 3 times in 1 mL PBS for 5 min at room temperature and coverslips were mounted on glass slides with Fluoromount-G.

In the case of samples prepared for STED microscopy, U20S Sec61ß-mEmerald cells were fixed with PBS-4% PFA (for 15 min, washed 3 times with 1mL PBS, and incubated with a permeabilization and blocking solution (PBS supplemented with 0.2% Triton X-100, 0.25% fish gelatin (Sigma Aldrich, G7765) 5% Normal Donkey serum (Merck, S30) and 3% Normal Goat Serum (Merck, S26)) for 1h at room temperature. Cells were then incubated overnight at 4°C with primary antibodies directed against NP and GFP to amplify the Sec61ß-mEmerald signal (Abcam 13970, 1:500) in staining buffer (PBS supplemented with 0.2% Triton X-100, 0.125% fish gelatin, 5% Normal Donkey serum and 3% Normal Goat Serum). Cells were then washed 3 times in 0.05% Tween 20-PBS and stained with an appropriate secondary antibody conjugated to an AF dye (AF594, ThermoFisher A-11042; Atto 647N, Merck 50185) in staining buffer for 1h at room temperature. Cells were then washed 3 times in 0.05% Tween 20-PBS, once with 1mL PBS and once with water. Coverslips were mounted on glass slides with Prolong Gold antifade reagent (Invitrogen, P10144).

#### Immunostaining for PI4P and PI3P

Cells were fixed with PBS-4% PFA (Fisher scientific, 47377) for 20 min, washed with PBS, and incubated for 20 min in blocking buffer (PBS with 5% bovine serum albumin, or PBS-5% BSA). For PI3P labeling, cells were incubated with purified FYVE-GST recombinant protein at a final concentration or 20 μg/mL in PBS with 0,5% saponin and 5% BSA for 1 h at room temperature, washed with PBS, and incubated with an anti-GST antibody (Rockland, 600-143-200, 1:300) in PBS-5% BSA for 1h at room temperature. For PI4P labeling, cells were incubated overnight at 4°C with an anti-PI4P antibody (Echelon-Z-P004, 1:150) in PBS with 0,5% saponin and 5% BSA, washed with PBS, incubated for 1h at room temperature with an appropriate secondary antibody conjugated to an Alexa Fluor dye (donkey anti-mouse IgG, Life Technologies), and post fixed 10 min with PBS-2% PFA.

#### Confocal microscopy

For confocal microscopy, Leica TCS SP8 or Nikon Confocal AX scanning confocal microscopes equipped with HC PL APO CS2 40X (NA=1.3) or 63X (NA=1.4) oil objectives were used. A MAX z-projection was applied to 5 to 10 stacks (0.35 µm step-size) in all figures. The fluorescence signals were acquired with the LAS X software (Leica) and analysed with Fiji (ImageJ) to determine fluorescence intensity profiles on a single focal plane. Cell Profiler [20,21] was used to determine the ratios of CLIMP63+ to RTN3+ areas. A SUM z-projection was applied to stacks of 15 images (0.35 µm step-size) and segmentation of the cells was performed by selecting the “Identify Primary Objects” module, using an adaptive, 2-class Otsu thresholding method. The number of PI4P punctae and their localization were manually analyzed using the “Cell counter” plugin in Fiji software. Line scans were recorded using the “Color Profiler” plugin (Color_Profiler. jar, https://imagej.nih.gov/ij/plugins/color-profiler.html) in Fiji software.

#### STED microscopy

STED images were acquired using theconfocal laser scanning microscope LEICA SP8 STED 3DX equipped with a 93×/1.3 NA glycerol immersion objective and with 3 hybrid detectors (HyDs). The specimens were excited with a pulsed white-light laser (598 nm or 640 nm) and depleted with a pulsed 775 nm depletion laser to acquire nanoscale imaging using SMD HyD detector. STED Images (2048 x2048 px) were averaged 16 times in line and acquired with a magnification zoom >2 leading to a pixel size in the range of 20-30nm. Excitation laser was adjusted to avoid any saturating pixels, and the same laser intensity and HyD sensitivity were used for both control and infected cells.

The levels of association of NP with Sec61ß were analyzed using a dedicated automatized program designed by L. Danglot using the Icy software [60] and “protocol” plugin. Pearson coefficient analysis was performed using the Colocalization Studio Icy Block [61] that takes into account both diffuse and/or aggregated signals. Cell area was automatically delineated using Easy Cell shape plugin (10.5281/zenodo.4317782) that rely on HK Means segmentation. Cell area was then converted to the Region of interest (Cell ROI) and Pearson coefficient was calculated with Sec61ß and NP channels within the Cell ROI from 18 control cells and 19 infected cells (sampled from 3 independent experiments). To analyse the Pearson coefficient in the surroundings of the nuclear membrane, we delineated a “perinuclear region” band of 100 pixels around the nucleus, contained within the Cell ROI.

#### Immunoblots

Total cell lysates prepared in RIPA or Laemmli buffer were loaded on 4-12% gradient acyrylamide gels (Invitrogen, NP0323). Immunoblot PVDF membranes (GE Healthcare Life science, Amersham Hybond 10600023) were incubated with HRP-conjugated streptavidin (Cell Signaling technology, 3999S, 1:2000) or with primary antibodies directed against RAB11 (Invitrogen 71-5300, 1:500), CLIMP63 (R&D systems AF7355, 1:500), RNT3 (Santa Cruz sc-374599, 1:500), ATG16L1 (MBL PM040, 1:1000), PB2 (GeneTex GTX125925, 1:5,000), Tubulin (Sigma-Aldrich, T5168 – B-5-1-2, 1:10,000), Histone-3 (Cell Signaling Technology, 9715S, 1:10,000) and revealed with appropriate secondary antibodies (Sigma Aldrich, A9044 and A9169, 1:10,000) and the ECL 2 substrate (Pierce). The chemiluminescence signals were acquired using the Chemidoc imaging system (Biorad) and a semi-quantitative analysis was performed with the ImageLab software (BioRad). Uncropped immunoblots are provided in **S4 File**.

#### siRNA-based assays

Small interfering RNAs (siRNAs) (Dharmacon ON-TARGETplus SMARTpools and Non-targeting Control pool) were purchased from Horizon Discovery, or from QIAGEN in the case of ATG16L1 siRNAs (Qiagen, SI04317418 and SI04999134). To assess the production of infectious viral particles, A549 cells seeded in 96-well plates (1.2×10^4^ cells/well) were transfected with 30 nM of siRNA, using 0.3 µL of the DharmaFECT1 transfection reagent (Horizon Discovery), and infected at 48 hours post-transfection (hpt) with the WSN virus at a MOI of 0.001 PFU/cell. Plaque assays were performed on MDCK cells as described in [62]. To assess subcellular features, A549 cells seeded in 24-well plates (7.5×10^4^ cells/well) were transfected with 30 nM of siRNA, using 3 µL of DharmaFECT1. At 24 hpt, cells were trypsinized, seeded on glass coverslips (13 mm diameter, #1.5, Epredia, CB00130RAC20MNZ0) in 24-well plates (10^5^ cells/well), and infected 24 h later with the WSN virus at a MOI of 5 PFU/cell. Immunostainings were performed as described above. Cell viability in the presence of siRNAs was assessed in the in 96-well plate format, using the CellTiter-Glo Luminescent Viability Assay kit (Promega). Knockdown efficiency of individual transcripts was quantified by RT-qPCR using SYBR green (ThermoFisher, 4309155) with the LightCycler^R^ 480 system (Roche), and the primers listed in **S5 File,** or by western-blot using antibodies directed against RAB11A or ATG16L1 as indicated above. RNA levels were normalised to GAPDH and analysed using the 2^-ΔΔCT^ method [63].

#### Statistical analysis of cell-based data

Statistical tests were performed using the GraphPad Prism (v10) software. For immunostaining data, unpaired Student’s t-test or two-way ANOVA were performed. For viral PFU data, two-way ANOVA was performed after log10 transformation of the data and Dunnett’s test was used for multiple comparisons with respect to the NT reference.

## Acknowledgements

We thank Federica Ferrentino and Ulrike Eggert (King’s College London, United Kingdom) for their very helpful advice regarding Cell Profiler-based analyses, and Maxime Chazal and Nolwenn Jouvenet (Institut Pasteur, Paris, France) for infection experiments with the Zika virus. We thank Olivier Schwartz, Blandine Monel and Julian Buchrieser (Institut Pasteur, Paris, France) for providing U2OS cells stably expressing Sec61ß-mEmerald, and Andres Merits (University of Tartu, Estonia) for providing the antibody against Zika virus NS3.

We gratefully acknowledge the technical and scientific support from the UtechS Photonic Bioimaging (Imagopole, C2RT, Institut Pasteur, supported by ANR-1N-INBS-04; Investments for the Future), the Necker SFR technical platforms and especially Meriem Garfa-Traoré at the Cell imaging facility, and the NeurImag Imaging core Facility team (part of IPNP, Inserm U1266 and Université Paris Cité, and member of the national infrastructure France-BioImaging supported by the French National Research Agency, ANR-10-INBS-04). We thank the Leducq foundation for supporting the acquisition of the Leica SP8 Confocal/STED 3DX microscope.

This work was funded by the Agence Nationale de la Recherche (ANR-21-CE11-0010-03 to NN and LD, ANR-10-LABX-62 to NN and MM, ANR-19-CE16-0012 to LD, ANR-17-CE140030-02, ANR-17-CE13-0015-003, ANR 22-CE14-0019 and ANR 18-CE14-0006 to EM, and ANR-21-CE35-0007 to MM), the Human Frontiers Science Program (HFSP RPG0040/2019 to NN), the Fondation pour la Recherche Médicale (FRM, « labellisation équipe » to EM), and the DIM 1Health to MM. JBB was supported by the HFSP RPG0040/2019 and the ANR-21-CE11-0010-03 grants. CA was supported by the ANR 22-CE14-0019 grant. JDG is a recipient of a doctoral fellowship from the French Ministry of Research/Université Paris-Cité and a 4th year PhD FRM scholarship (grant FDT202304016558).

## Supporting Information

**S1 Fig.**
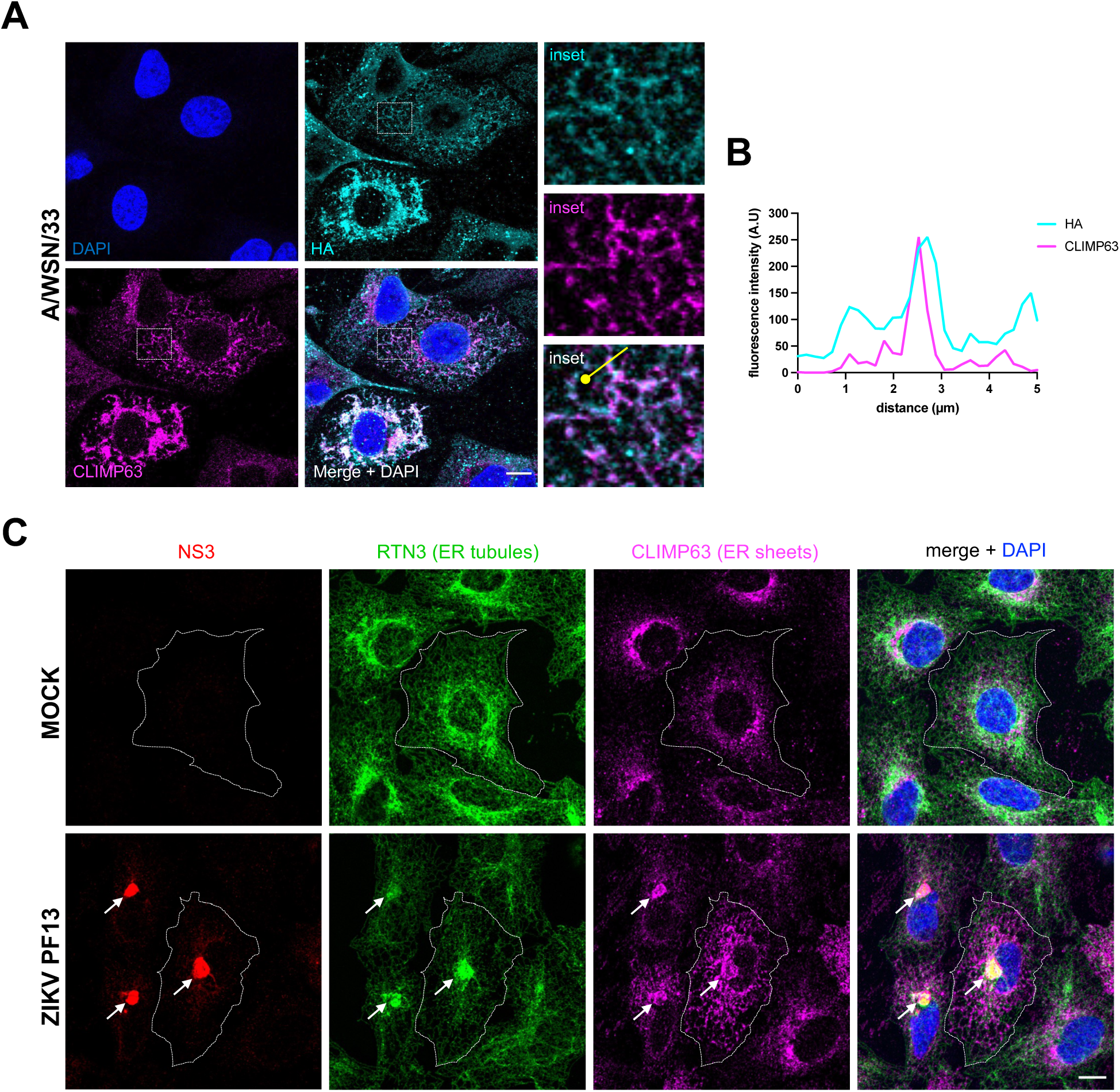
Viral-induced remodelling of the ER (related to Figure 1). **A.** A549 cells were infected with WSN at a MOI of 5 PFU/cell for 8 h. Fixed cells were stained for the viral HA and cellular CLIMP63 proteins. Nuclei were stained with DAPI and cells were imaged with a confocal microscope. Scale bar: 10µm. **B.** Fluorescence intensity profile for HA (cyan) and CLIMP63 (magenta) along the white line drawn in panel A (merge inset), starting from the knob. **C.** A549 cells were infected with ZIKV PF13 at a MOI of 5 PFU/cell for 24 h, or mock-infected. Fixed cells were stained for the viral NP and the cellular RTN3 and CLIMP63 proteins. The RTN3 staining was used to delineate the cell edges. Nuclei were stained with DAPI and cells were imaged with a confocal microscope. White arrows indicate viral factories surrounded with remodelled ER membranes. Scale bar: 10µm.

**S2 Fig.**
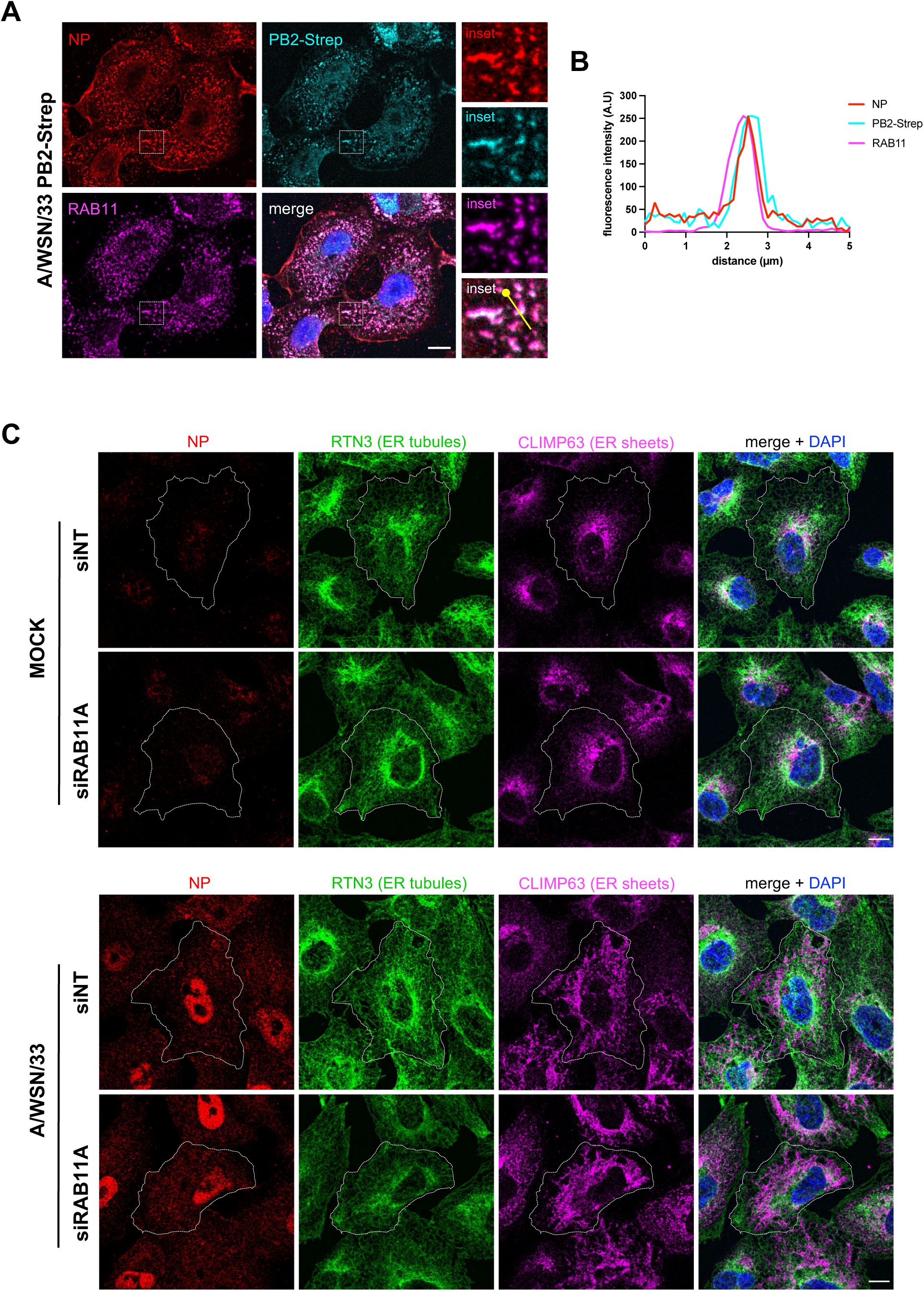
Impact of RAB11A depletion on ER remodelling and vRNP localisation upon infection (related to Figure 2). **A.** A549cells were infected with the WSN-PB2-Strep virus at a MOI 5 PFU/cell for 8 h. Fixed cells were stained for NP, PB2 (StrepTactin-488) and RAB11. Nuclei were stained with DAPI and cells were imaged with a confocal microscope. Scale bar: 10µm. **B.** Fluorescence intensity profile for NP (red), PB2-Strep-tag (cyan) and RAB11 (magenta) along the white line drawn in panel A (merge inset), starting from the knob. **C.** A549 cells were treated with RAB11A-specific or control Non-Target (NT) siRNAs for 48 h, and subsequently infected with WSN at a MOI of 5 PFU/cell for 8 h, or mock infected. Fixed cells were stained for the viral NP and cellular RTN3 and CLIMP63 proteins. The difference in permeabilisation protocols (saponin versus Triton) most likely accounts for the difference in NP signal patterns in this experiment compared to the experiment shown in Figure 2A. Scale bar: 10µm.

**S3 Fig.**
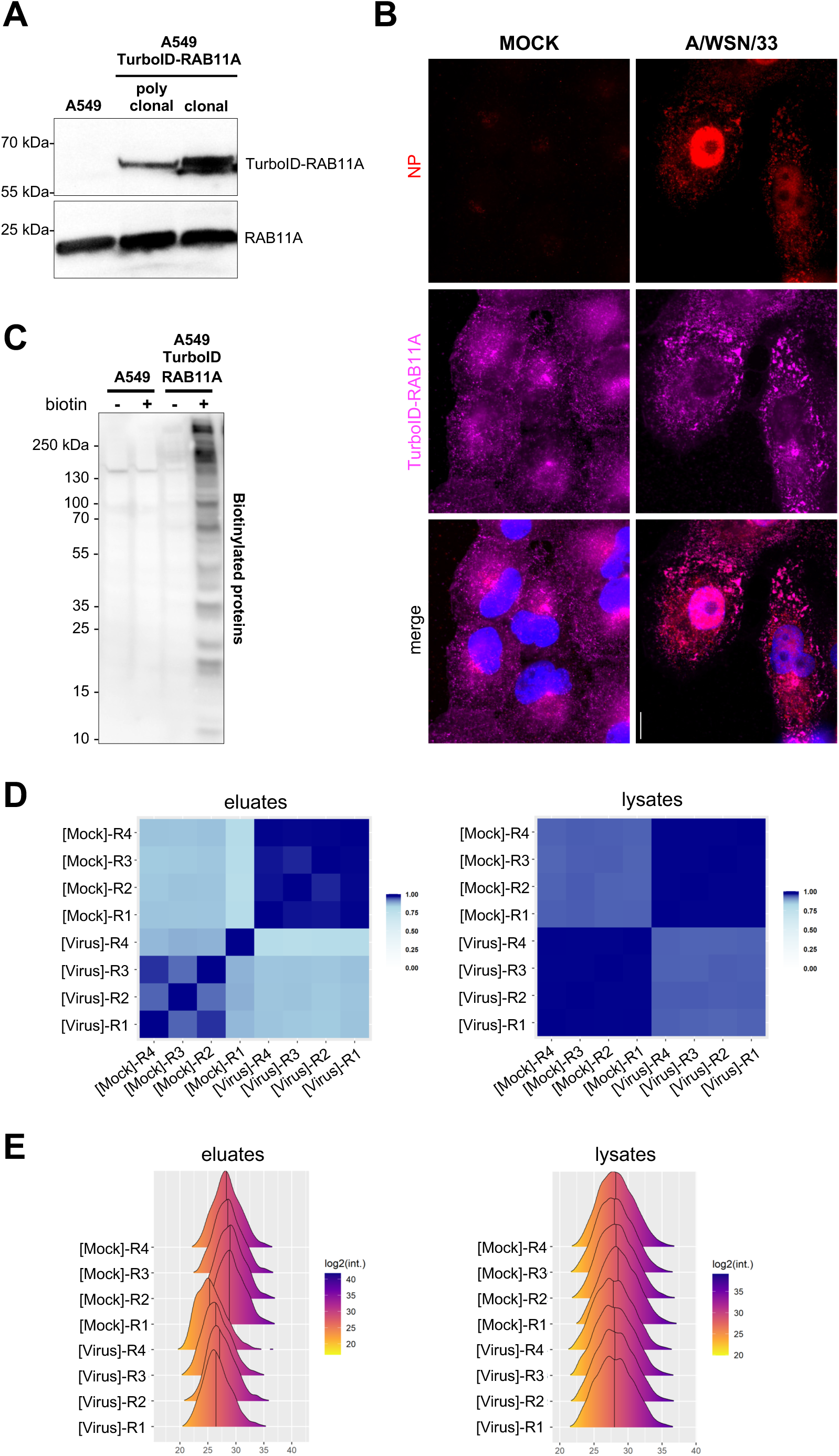
RAB11A proximity labelling (related to Figure 3). **A-C.** Characterization of the clonal population of A549cells stably expressing TurboID-RAB11A. (A) Total cell lysates of A549 parental cells (untransduced), a polyclonal population of transduced A549-TurboID-RAB11A cells, and the clonal population isolated thererof, were analysed by western-blot using antibodies specific for RAB11. (B) The clonal A549-TurboID-RAB11A cells were infected with the WSN virus at a MOI of 5 FPU/cell for 8 h. Fixed cells were stained for the HA-tag (3xHA-TurboID-RAB11A) and the viral NP. Nuclei were stained with DAPI and cells were imaged with a confocal microscope. Scale bar: 10µm. (C) Total cell lysates of parental A549 cells or clonal A549-TurboID-RAB11A cells, incubated or not for 10 mn at 37°C in the presence of 50 µM biotin, were analysed by western-blot using Strep-Tactin conjugated with HRP to detect biotinylated proteins. **D.** Correlation matrices between replicates of eluates (left) and total lysates (right). A correlation matrix represents the Pearson correlation coefficients between each pair of samples computed using all complete pairs of intensity values measured in these samples. Intensity values correspond to TMT-MS2 quantitative relative abundance metrics in the columns titled “Reporter intensity corrected” of the “proteinGroups.txt” file of MaxQuant. The samples identification numbers are indicated in the format “Mock.number of the technical replicate” and “IAV.number of the technical replicate”. Pearson correlation coefficients are indicated in the lower triangular parts of the matrices. In the upper triangular parts, the diameters and gradient colors of the circles are function of these coefficients. **E.** Distributions of the log2(intensities) for the proteins without missing values in the eluates (left) and total lysates (right). The samples identification numbers are indicated on the vertical axis in the format “[Mock].number of the technical replicate” and “[Virus].number of the technical replicate”.

**S4 Fig.**
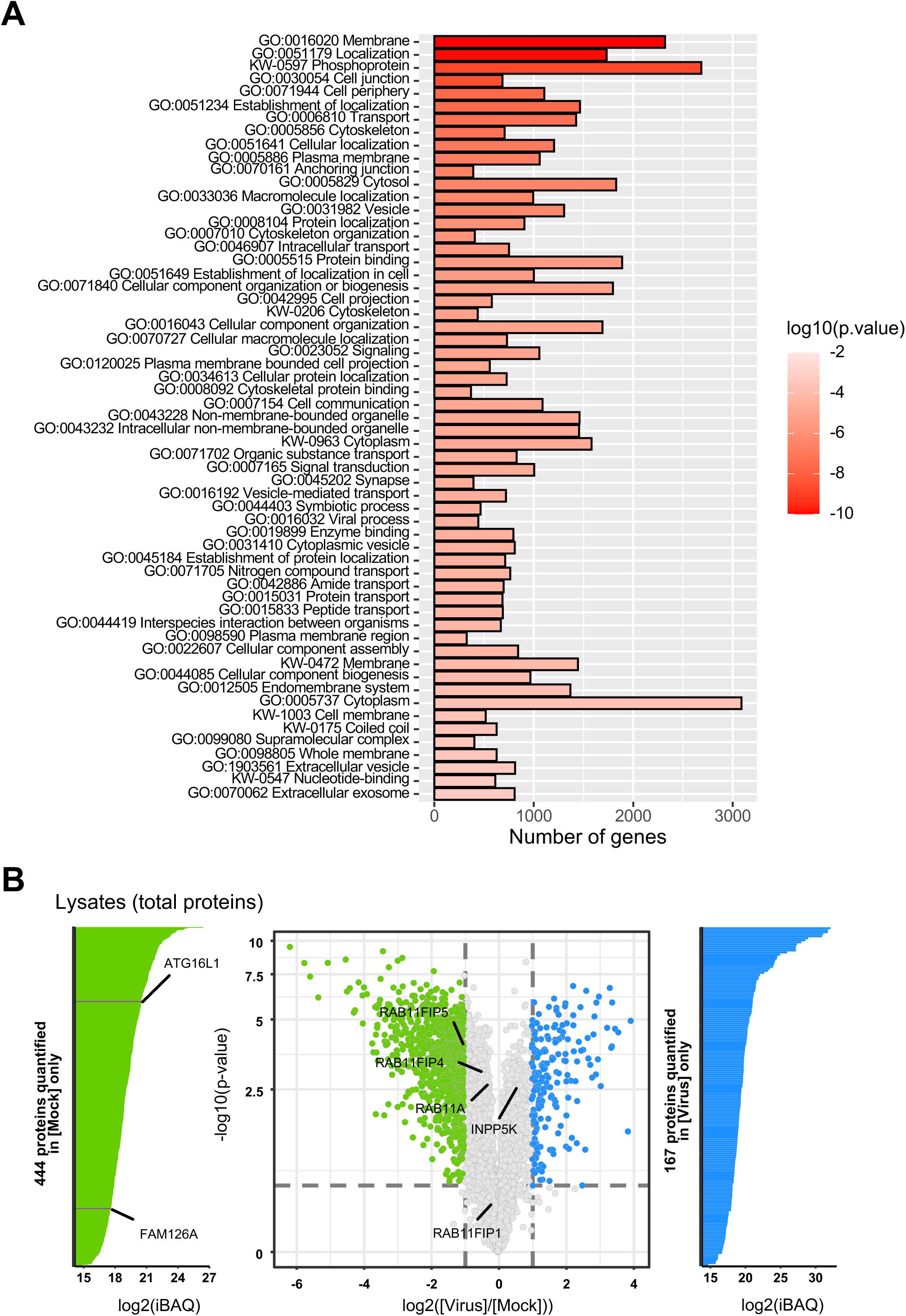
RAB11A proximity labelling (related to Figure 3) **A.**Gene Ontology (GO) term enrichment analysis of the set of 3775 biotinylated proteins identified as being enriched in the biotynylated samples across all conditions and replicates, over a total of 6011 proteins identified in biotinylated eluates and/or total lysates across all conditions and replicates (both sets of proteins are listed in the Supp File 1). The graph represents the number of genes corresponding to each indicated category (x axis) and the enrichment p-value (color scale). **B.** Volcano plot showing the log2 fold change (x axis) and its significance (-log10(p-value), y axis) associated to a False Discovery Rate < 1%) for each protein (dots) in total lysates from the RAB11A proximity labelling experiment. The log2 fold change refers to the enrichment in WSN-infected (n=4) versus mock-infected (n=4) samples. Blue and green dots represent proteins enriched in WSN-infected versus mock-infected samples, and proteins enriched in mock-infected versus WSN-infected samples, respectively. The iBAQ plots are shown on the sides of the volcano plot for proteins that are present in WSN-infected samples only (blue) or in mock-infected samples only (green). E and G. A549 cells were treated with control non-target (NT) siRNAs or with siRNAs targeting FAM126A, INPP5K (E) or ATG16L1 (G) for 48 h, and subsequently infected with WSN at a MOI of 5 PFU/cell for 8 h, or mock-infected. Fixed cells were stained for PI3P or PI4P. Nuclei were stained with DAPI and cells were imaged with a confocal microscope. Scale bar: 10 µm **F and H**. A549 cells treated as in E and G, respectively were analyzed with the Fiji software to determine the mean intensity of the PI3P or PI4P signal per cell. Each dot represents one cell, and the data from three independent experiments are shown (black, grey and blue dots). The median and standard deviation values are represented (100-150 cells per condition). **\*\*\*** : p-value <0.001, **\*\*\*\*** : p-value <0.0001, ns:non significant, unpaired t-test.

**S5 Fig.**
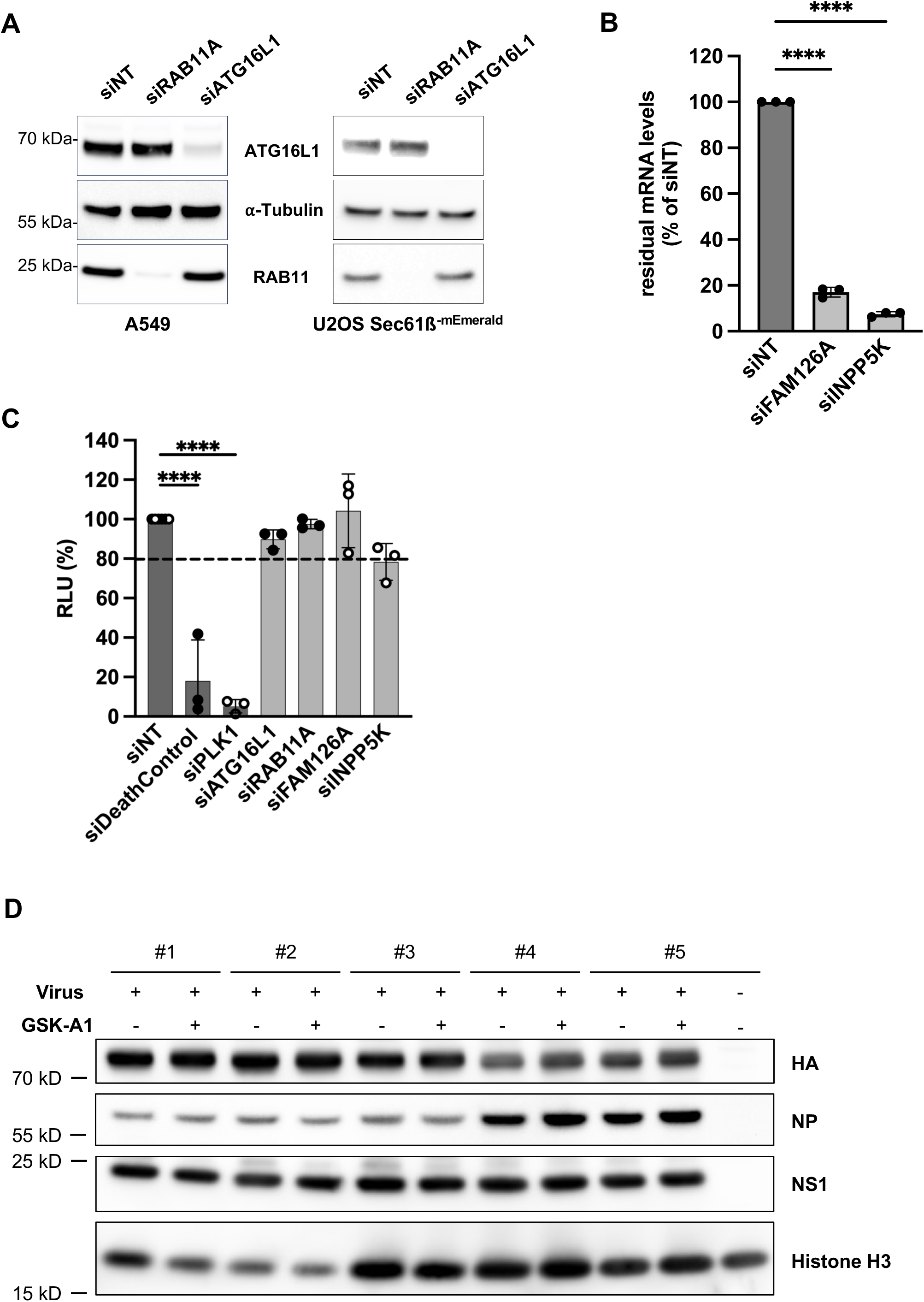
SiRNA-mediated knock-down efficiency and cell viability assessments. **A-B.** Knock-down efficiency of siRNA pools. A549 cells were treated with the indicated siRNAs for 48 hours. (A) Total cell lysates were prepared and analysed by western blot, using the indicated antibodies. (B) Total RNA were extracted and analysed by RTqPCR using gene specific primers. The residual mRNA levels are expressed as percentages (100%: NT siRNA). Data shown are the mean ± SD of three experiments performed in triplicates. ****p <0.0001 (one-way ANOVA and Dunnett’s multiple comparisons test, reference: NT siRNA). **C.** Cell viability upon treatment with siRNA pools. A549 cells were treated with the indicated siRNAs for 48 hours and cell viability was determined at 72 hpt using the CellTiter-Glo Luminescent Viability Assay (Promega). The data shown (RLU: Relative Light Units) are expressed as percentages (100%: NT siRNA) and are the mean ± SD of three independent experiments performed in triplicates. Black and white dots correspond to two distinct series of experiments, in which the “Death Control siRNA” (xxx) and a siRNA directed against PLK1 were used as positive controls, respectively. The dotted line indicates a 20% reduction in luciferase signal. ****p<0.0001 (one-way ANOVA and Dunnett’s multiple comparisons test, reference: NT siRNA, no indication means no significant difference). **D.** A549 cells were infected with WSN at a MOI of 5 PFU/cell. Two hours later, the GSK-A1 drug was added at a final concentration of 100 nM. At 6 hpi, total cell lysates from five independent experiments (#1 to #5) were prepared and analysed by western blot, using the indicated antibodies. Cropped blots are shown.

**S1_File.** Proteins detected by mass spectrometry in the lysates and eluates

**S2_File.** Differential analysis of mass spectrometry data from eluates

**S3_File.** Differential analysis of of mass spectrometry data from lysates

**S4_File.** Uncropped western blot membranes

**S5_File.** Primers used for RTqPCR

